# 3D Mitochondrial Structure in Aging Human Skeletal Muscle: Insights into MFN-2 Mediated Changes

**DOI:** 10.1101/2023.11.13.566502

**Authors:** Estevão Scudese, Zer Vue, Prassana Katti, Andrea G. Marshall, Mert Demirci, Larry Vang, Edgar Garza López, Kit Neikirk, Bryanna Shao, Han Le, Dominique Stephens, Duane D. Hall, Rahmati Rostami, Taylor Rodman, Kinuthia Kabugi, Chanel Harris, Jian-qiang Shao, Margaret Mungai, Salma T. AshShareef, Innes Hicsasmaz, Sasha Manus, Celestine Wanjalla, Aaron Whiteside, Revathi Dasari, Clintoria Williams, Steven M. Damo, Jennifer A. Gaddy, Brian Glancy, Estélio Henrique Martin Dantas, André Kinder, Ashlesha Kadam, Dhanendra Tomar, Fabiana Scartoni, Matheus Baffi, Melanie R. McReynolds, Mark A. Phillips, Anthonya Cooper, Sandra A. Murray, Anita M. Quintana, Vernat Exil, Annet Kirabo, Bret C. Mobley, Antentor Hinton

## Abstract

Age-related atrophy of skeletal muscle, is characterized by loss of mass, strength, endurance, and oxidative capacity during aging. Notably, bioenergetics and protein turnover studies have shown that mitochondria mediate this decline in function. Although exercise has been the only therapy to mitigate sarcopenia, the mechanisms that govern how exercise serves to promote healthy muscle aging are unclear. Mitochondrial aging is associated with decreased mitochondrial capacity, so we sought to investigate how aging affects mitochondrial structure and potential age-related regulators. Specifically, the three-dimensional (3D) mitochondrial structure associated with morphological changes in skeletal muscle during aging requires further elucidation. We hypothesized that aging causes structural remodeling of mitochondrial 3D architecture representative of dysfunction, and this effect is mitigated by exercise. We used serial block-face scanning electron microscopy to image human skeletal tissue samples, followed by manual contour tracing using Amira software for 3D reconstruction and subsequent analysis of mitochondria. We then applied a rigorous *in vitro* and *in vivo* exercise regimen during aging. Across 5 human cohorts, we correlate differences in magnetic resonance imaging, mitochondria 3D structure, exercise parameters, and plasma immune markers between young (under 50 years) and old (over 50 years) individuals. We found that mitochondria we less spherical and more complex, indicating age-related declines in contact site capacity. Additionally, aged samples showed a larger volume phenotype in both female and male humans, indicating potential mitochondrial swelling. Concomitantly, muscle area, exercise capacity, and mitochondrial dynamic proteins showed age-related losses. Exercise stimulation restored mitofusin 2 (MFN2), one such of these mitochondrial dynamic proteins, which we show is required for the integrity of mitochondrial structure. Furthermore, we show that this pathway is evolutionarily conserved as Marf, the MFN2 ortholog in *Drosophila*, knockdown alters mitochondrial morphology and leads to the downregulation of genes regulating mitochondrial processes. Our results define age-related structural changes in mitochondria and further suggest that exercise may mitigate age-related structural decline through modulation of mitofusin 2.

**Graphical Abstract:**
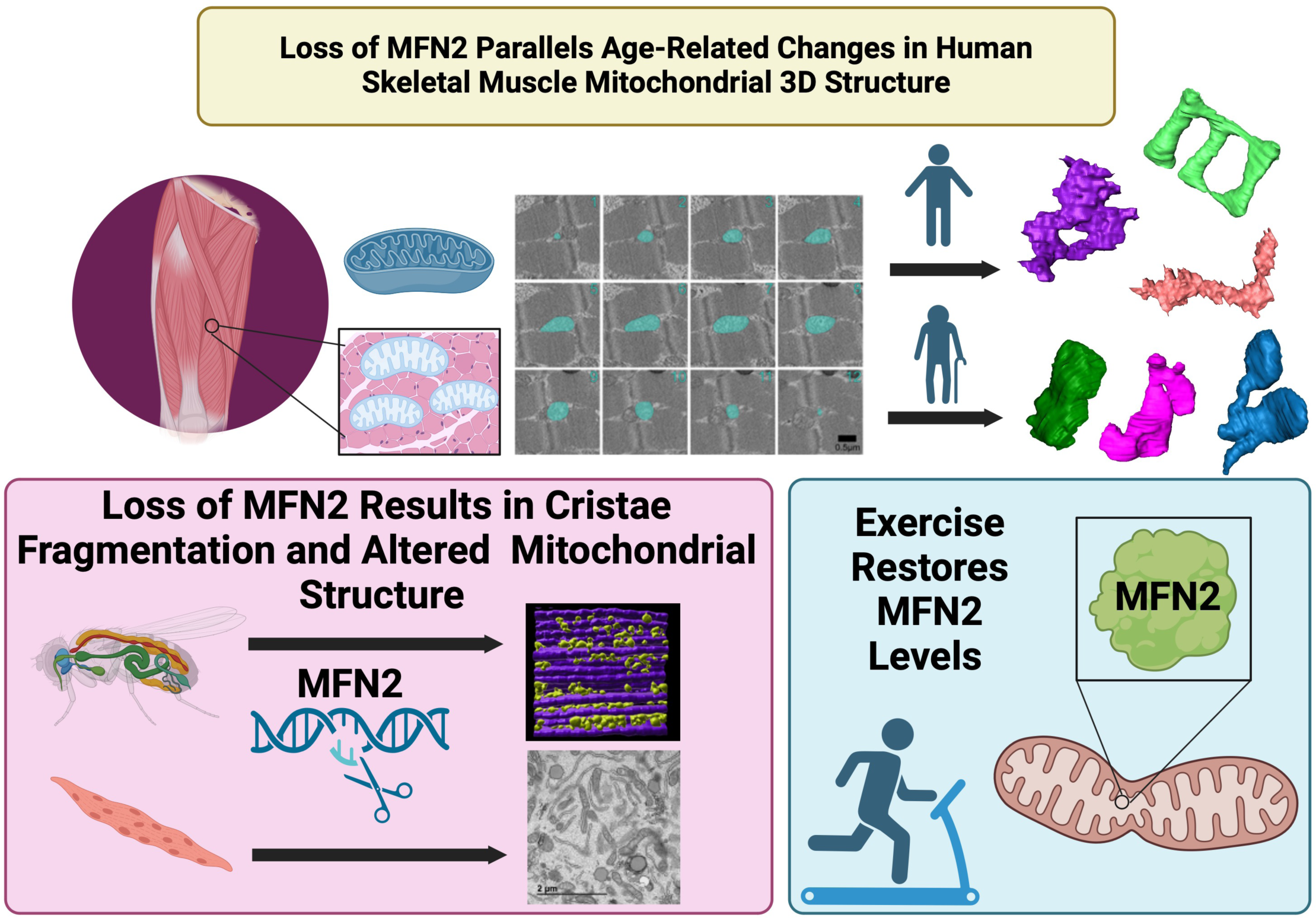
Age-related skeletal muscle atrophy shows morphological alterations in mitochondrial structure associated with declining function. Our findings propose that exercise intervention may counteract these structural declines by reinstating levels of mitofusin 2, thus highlighting a potential mechanism by which exercise attenuates age-induced mitochondrial dysfunction.

## Introduction

Aging is an inescapable biological process characterized by a progressive decline in physiological and metabolic functions. Across this process, changes in metabolic homeostasis, muscle mass, and function have been observed across species and sexes (Greenlund & Nair 2003; Tay et al. 2015). The pluralistic effects of aging contribute to an increasing vulnerability to disease, particularly in skeletal muscle. Sarcopenia, which consists of muscle degeneration and progressive loss of skeletal muscle mass, remains a global health issue (Martinez et al. 2015; Greenlund & Nair 2003). While the prevalence of sarcopenia remains poorly elucidated, past cross-sectional analyses have shown that approximately 20% of hospitalized adult patients meet the criteria for sarcopenia, with risk factors including old age and obesity (Sousa et al. 2015). Currently, sarcopenia accounts for approximately 1.5% of healthcare expenditures in the United States, and as the population worldwide ages, the burden of sarcopenia will only be exacerbated (Coen et al. 2019; Filippin et al. 2015). Generally, sarcopenia occurs at approximately 40 years of age with a loss of 8% of muscle mass per decade; then, the rate increases to 15% of muscle mass loss per decade after 70 years (Kim & Choi 2013). Sarcopenia, in turn, increases the risk of mortality and disability by exacerbating the risk of adverse events including falls, fractures, and functional decline, cumulatively resulting in a decreased quality of life (Filippin et al. 2015). There is limited understanding of how changes in mitochondria during the aging process contribute to muscle dysfunction and the development of age-related characteristics.

Aging in skeletal muscle has been studied to reduce its burden on healthcare systems globally, yet therapies remain limited. No clinical treatment, other than nutritional changes and a regular exercise regimen, exists to mitigate sarcopenia development (Phu et al. 2015; Taaffe 2020). Still, the underlying molecular mechanisms driving these age-related changes and the reason why exercise may be able to reverse them in some disease states remains insufficiently understood. Recently, mitochondria have emerged as a potential target for the mitigation of age-related atrophy because mitochondrial functional changes often precede hallmarks of sarcopenia: the loss of muscle mass and function (Campo et al. 2018; Coen et al. 2019; Hepple 2014). Although few studies examine the therapeutic value of exercise with aging independent of sarcopenia, recent findings have shown that regular exercise, through mitochondrial-dependent mechanisms, counteracts the deleterious effects of aging in skeletal muscle (Grevendonk et al. 2021). The role of mitochondria in health is clear: in *Caenorhabditis elegans,* mitochondrial content correlates strongly with lifespan, with mitochondrial networking declining antecedent to sarcomere loss (Gaffney et al. 2018; Schriner et al. 2005). In addition to the process of aging, mitochondrial dysfunction has been shown in the skeletal muscle of individuals with a range of medical conditions, including chronic kidney disease, congestive heart failure, and diabetes (Kim et al. 2008; Gamboa et al. 2016; Scandalis et al. 2023). The high prevalence of sarcopenia and physical dysfunction among these individuals underscores the importance of muscle health and underlying mitochondria dysfunction. We have previously shown, in a murine model, that mitochondrial structure in skeletal muscle undergoes reductions in size and changes in morphology during aging, which may confer reduced functional capacity prior to the development of sarcopenia (Vue, Garza-Lopez, et al. 2023). Together, these findings suggest mitochondrial structure may precede sarcopenia, but structural changes in human skeletal muscle during general age-related atrophy remain poorly defined, especially in relation to exercise.

While mitochondria are generally characterized by their role in ATP synthesis, their pluralistic roles extend far beyond this, including apoptosis, cellular metabolic and redox signaling and calcium homeostasis, cumulatively linking mitochondria to the aging process (Bratic & Larsson 2013; Campo et al. 2018; Jenkins et al. 2024). These organelles are not static but highly dynamic, undergoing constant cycles of fission, mediated by effectors such as dynamin-related protein 1 (DRP1), and fusion, mediated by effectors such as optic atrophy protein 1 (OPA1) and mitofusins 1 and 2 (MFN1 and MFN2), to adapt to the cellular environment (Dong et al. 2022; Chan 2012). Interruptions of either of these processes can interfere with mitochondrial function, cause dysfunction, and be representative of pathology (Chan 2012; Bartsakoulia et al. 2018; Chen et al. 2005). Recently, an age-associated loss of OPA1 was linked to reduced skeletal muscle mass (Tezze et al. 2017). Furthermore, OPA1, when increased by exercise, can increase mitochondrial calcium uniporter-dependent mitochondrial Ca^2+^ uptake (Zampieri et al. 2016). Of particular interest in age-related atrophy is MFN2, since age-related losses of MFN2 have been shown to underlie metabolic alterations and sarcopenia (Sebastián et al. 2016). More recently, overexpression of MFN2 in skeletal muscles of young and old mice has been shown to cause mild non-pathological hypertrophy and potentially mitigate aging-related muscle atrophy (Cefis et al. 2024).

Beyond fusion and fission dynamics, these same regulators may often give rise to alterations in the mitochondrial network and morphology (Chen et al. 2005; Liu & Hajnóczky 2011). Thus, it is understood that mitochondria change their three-dimensional (3D) morphology to unique phenotypes that are representative of their cellular state, such as during oxidative stress (Glancy et al. 2020). For instance, donut-shaped mitochondria may emerge as a pathology-induced mechanism to increase surface area at the expense of volume (Hara et al. 2014). Decreased volume results in less space for the folds of the inner mitochondrial membrane, known as cristae, which optimize ATP synthesis (Cogliati et al. 2016). However, the increased surface area may also allow for increases in mitochondrial–endoplasmic reticulum contact sites (MERCs), which may function in alternative biochemical roles such as in calcium homeostasis (Bustos et al. 2017). In tandem, alterations in mitochondrial dynamics can result in unique 3D structures that may play pivotal roles in the aging process and the associated muscle dysfunction.

Previously, we performed 3D reconstruction for the volumetric rendering of mitochondria using manual contour tracing, which provides information on mitochondrial phenotypes, including those in murine skeletal muscle during aging (Vue, Garza-Lopez, et al. 2023). This method is facilitated by serial block-face scanning electron microscopy (SBF-SEM), which given its large range, allows for large-volume renderings and mitochondrial networks to be accurately replotted (Marshall, Neikirk, et al. 2023; Courson et al. 2021). Other studies have examined the 3D structure of human skeletal muscle in the context of mitochondrial DNA (mtDNA) diseases (Vincent et al. 2019). Yet, in the context of aging, human skeletal muscle changes remain poorly elucidated. Traditional 2D techniques examining aged skeletal muscle cells of humans have shown large mitochondria with disrupted cristae (Beregi et al. 1988). However, mitochondria in aged human skeletal muscle tissue were observed to shrink in a sex-dependent manner (Callahan et al. 2014), and aged mouse skeletal muscle tissue showed increased branching during aging due to an increased MFN2-DRP1 ratio (Leduc-Gaudet et al. 2015). These conflicting results limit our understanding of mitochondrial ultrastructure in human skeletal muscle aging. To our knowledge, the 3D structure of human skeletal muscle mitochondria during the aging process has yet to be defined.

In this study, we employ a multi-pronged approach to explicate the interplay among aging, mitochondrial dynamics, and exercise therapies. Utilizing both human and murine models, as well as the genetically tractable model organism *Drosophila melanogaster*, we offer an analysis of how aging affects skeletal muscle mass and mitochondrial 3D architecture. We further investigate whether these age-related changes are evolutionarily conserved and explore the potential for exercise to mitigate these detrimental effects. Our findings offer insights into the role of mitochondrial structural changes in age-associated dysfunction and metabolic shifts. We also establish potential mechanisms for exercise as a modulatory tool for age-related deficiencies. We further highlight hematological changes in the aging process and show that mitochondrial structural rearrangement is mechanistically responsible for some of the therapeutic benefits that exercise exerts in the context of aging and age-related diseases. Through using 5 human cohorts (Supplemental Figure 1), we broadly correlate that in old individuals (50 years or more), as compared with younger individuals, magnetic resonance imaging shows gross skeletal muscle morphological changes, mitochondrial ultrastructure is reconfigured, muscle strength weakens, and blood biomolecules are altered. Our findings highlight MFN2 as a potential therapy for age-dependent skeletal muscle change in mitochondrial structure, suggesting a future mechanistic target for an effector that mediates exercise-dependent impairment of muscle atrophy.

## Methods

### Human Sample Cohort

Several different human cohorts across multiple countries were utilized for this study. Specimens for all 3D reconstructions (Figures 2–3) were collected at Vanderbilt University Medical Center. Collection of human quadriceps tissue was approved by the Vanderbilt University Institutional Review Board (IRB) under the title “Mitochondria in Aging and Disease--study of archived and autopsy tissue” with an associated IRB number of 231584. All other human samples were obtained from Brazilian cohorts according to the CAEE (Ethics Appreciation Presentation Certificate) guidelines. Samples from young individuals were collected, and experiments were performed under CAEE number 61743916.9.0000.5281; samples from older individuals were collected under CAEE number 10429819.6.0000.5285. Mixtures of male and female samples (specified in figures) were used in all studies, with a general cutoff age of ∼50 years for humans.

**Figure 1.**
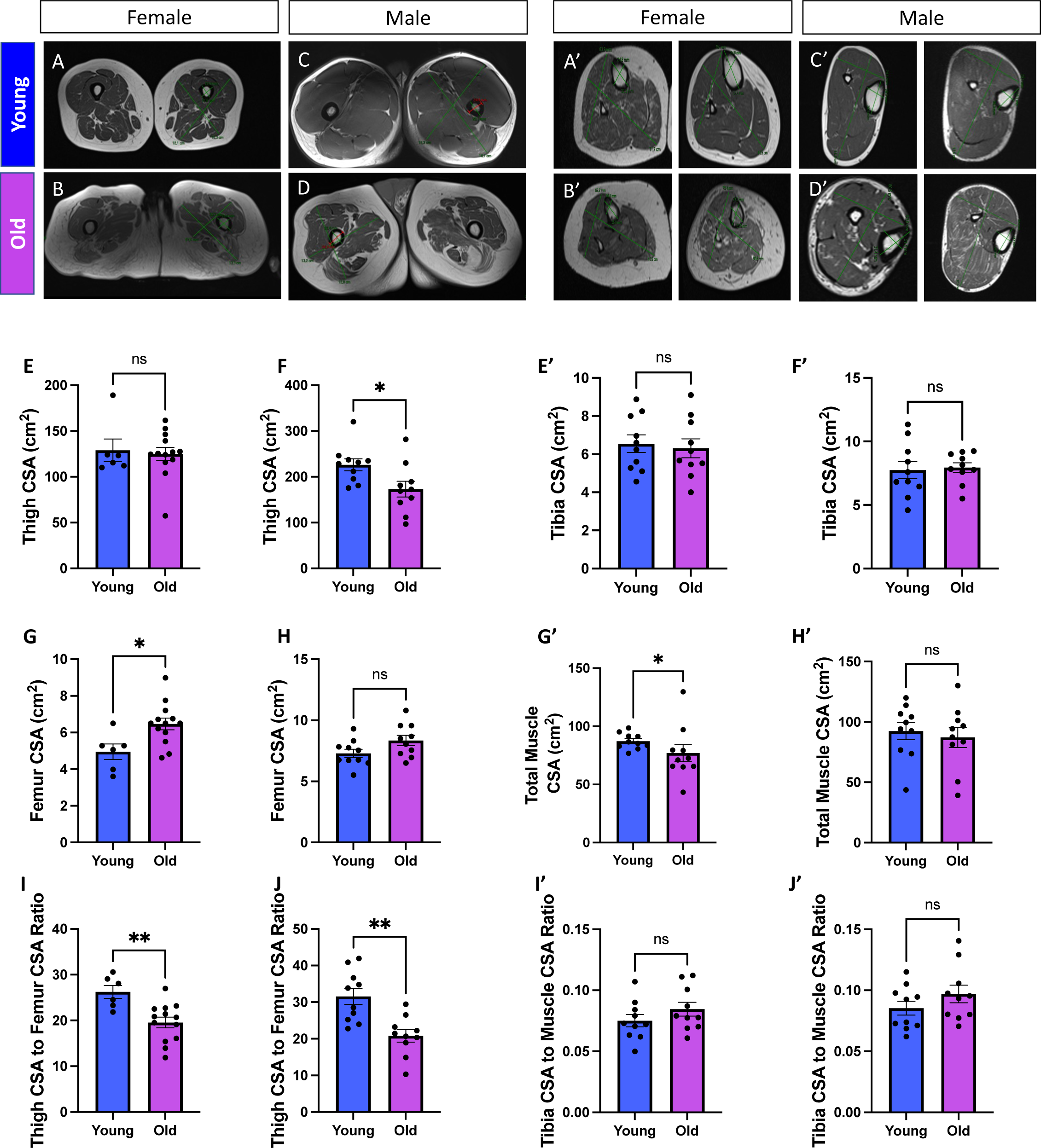
Comparative Analyses of Musculoskeletal Characteristics in Young and Old Participants Differentiated by Sex. **(A-G)** Cross-sectional imaging of thigh musculature and skeletal anatomy data from (A) females under 50 years old (aged 8–41 years old; n = 7), (B) females over 50 years old (aged 57–79 years old; n = 13), (C) males under 50 years old (aged 19–48 years old; n = 10), and (D) males over 50 years old (aged 67–85 years old; n = 10). (E) Thigh cross-sectional area (CSA) measurements for females and (F) males, with data for young and old participants represented by blue and purple bars, respectively. (G) Femur CSA measurements for females and (H) males. (I) Ratio of thigh CSA and femur CSA for females and (J) males. Individual data points indicating separate individuals (Supplemental File 1) are represented by dots on the bar graphs. (A’-G’) Cross-sectional imaging of calf musculature and skeletal anatomy data from (A’) females under 50 years old (aged 15–48 years old; n = 10), (B’) females over 50 years old (aged 54–82 years old; n = 10), (C’) males under 50 years old (aged 22–49 years old; n = 10), and (D’) males over 50 years old (aged 51–81 years old; n = 10). (E’) Tibia cross-sectional area (CSA) measurements for females and (F’) males, with data for young and old participants represented by blue and purple bars, respectively. (G’) Total muscle of calf CSA measurements for females and (H’) males. (I’) Ratio of tibia CSA to calf CSA for females and (J’) males. Individual data points indicating separate individuals (Supplemental File 2) are represented by dots on the bar graphs. Mann–Whitney tests were used for statistical analysis. Statistical significance is denoted as ns (not significant), *p < 0.05, and **p < 0.01.

**Figure 2.**
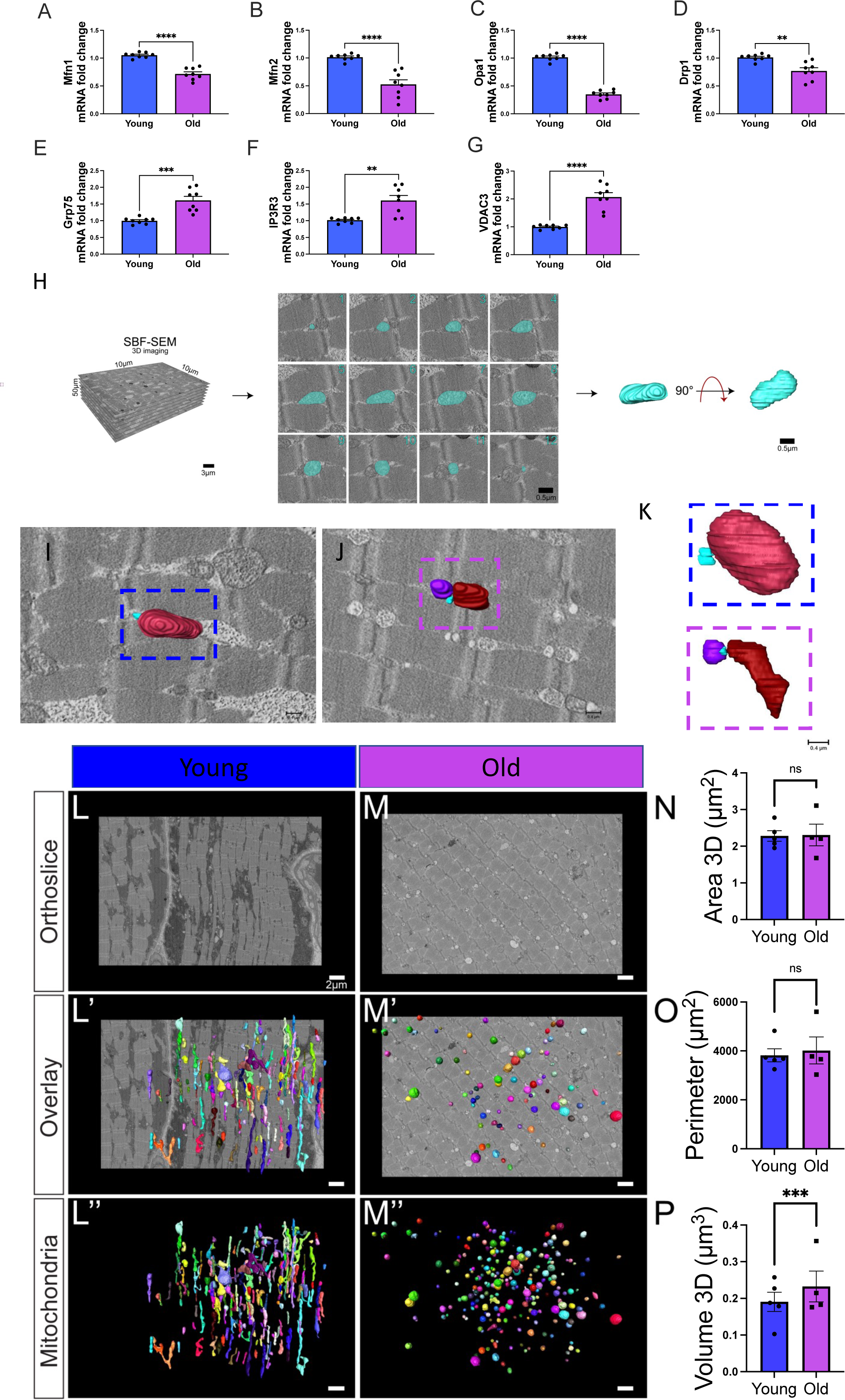
Changes in Mitochondrial Dynamics and Structure with Aging in Human Skeletal Muscle. (A–D) Quantified differences in mRNA fold changes, as determined by quantitative PCR, of various mitochondrial dynamic proteins and (E–G) mitochondrial–endoplasmic reticulum contact site proteins. Parameters are compared between the young and old groups (n=8 for both). (H) Workflow of serial block-face scanning electron microscopy (SBF-SEM) manual contour reconstruction to recreate 3D mitochondrial structure from young and old human samples. The workflow depicts SBF-SEM, allowing for orthoslice alignment, subsequent manual segmentation of orthoslices, and ultimately, 3D reconstructions of mitochondria. (I) Qualitative image of mitochondrial–endoplasmic reticulum contact sites in young and (J) old cohorts, with (K) specific contact sites magnified for viewing. Blue structures represent the endoplasmic reticulum. (L) Differences in orthoslice mitochondrial structure between young and (M) old human skeletal muscle, with a scale bar of 2 μm. (L’) Overlaid view of the segmented mitochondria on the orthoslice, emphasizing distinct mitochondrial shapes and distributions observed with 3D reconstruction in young and (M’) older participants. (L’’) 3D reconstructed images of isolated mitochondria from young and (M’’) older participants. (N) Differences in mitochondrial area, (O) mitochondrial perimeter, and (P) volume between the young and old groups. (A–G) Each dot represents an independent experimental run or (N-P) average of all mitochondria quantifications in each patient. 5 young individuals surveyed (mitochondrial number varies; Case #1: n = 253; Case #2: n = 250; Case #3: n = 250; Case #4: n = 252; Case #5: n = 253; total mitochondria surveyed across young cohort: n = 1258) and 4 old cases (mitochondrial number varies; Case #1: n = 254; Case #2: n = 250; Case #3: n = 250; Case #4: n = 250; total mitochondria surveyed across old cohort: n = 1004) for 3D reconstruction. Significance was determined with the Mann–Whitney test comparing the combined number of mitochondria in young (n = 1258) and old (n = 1004) cohorts, with ns, *, **, ***, and **** representing not significant, p ≤ 0.05, p ≤ 0.01, p ≤ 0.001, and p ≤ 0.0001.

**Figure 3.**
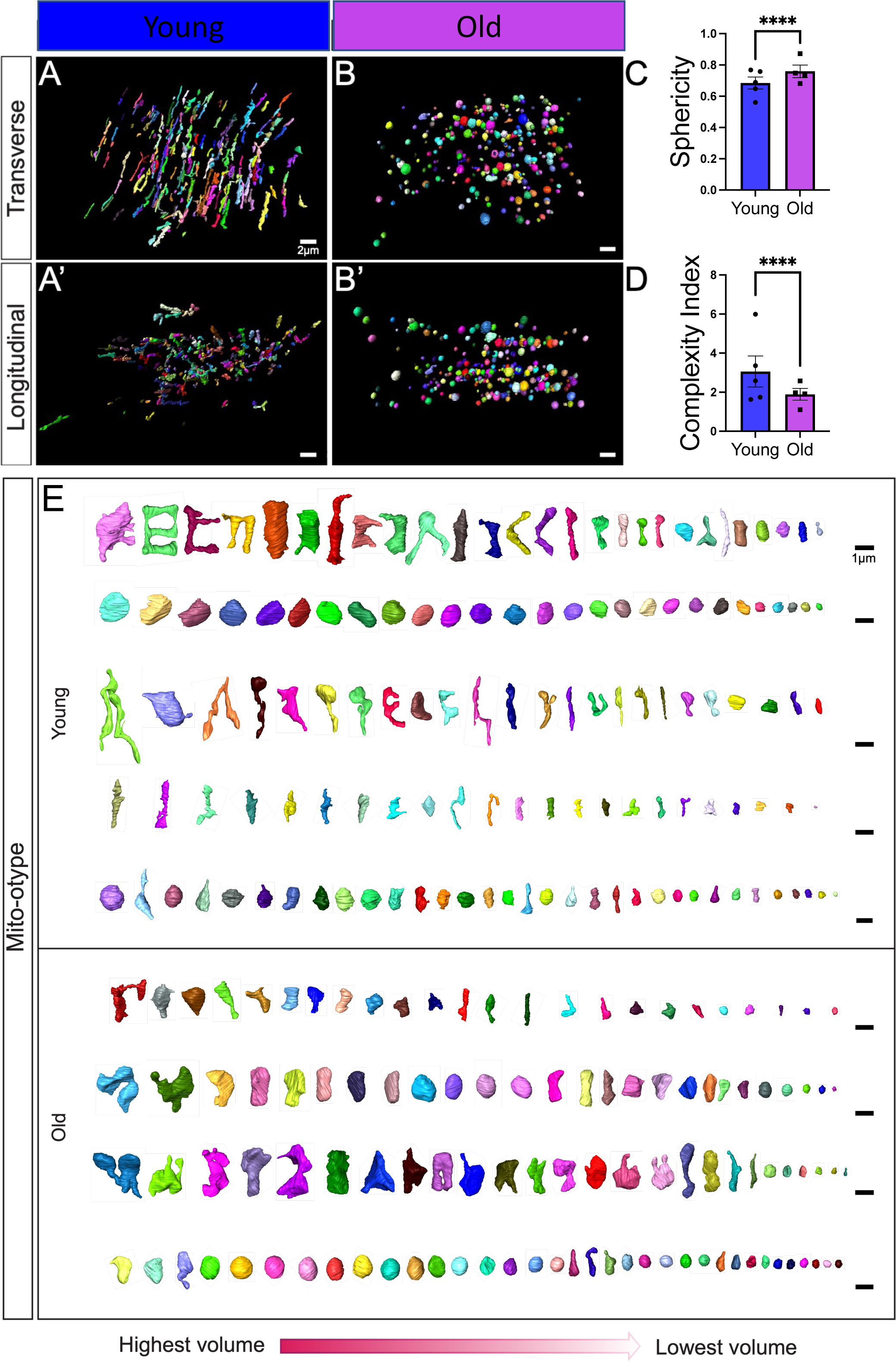
Changes in Mitochondrial Branching and Networking after Aging Revealed by Serial Block-face Scanning Electron Microscopy. (A) 3D reconstructions showing young and (B) old human skeletal muscle from a transverse point of view. (A’) 3D reconstructions showing young and (B’) old human skeletal muscle from a longitudinal point of view. (C) Sphericity of mitochondria in the young and old groups. (D) The mitochondrial complexity index (MCI), which is analogous to sphericity, was used to compare the young and old groups. (E) Mito-otyping was used to display the diversity of mitochondrial phenotypes, as ordered by volume, to show the mitochondrial distribution in the young and old groups, with each row representing an independent patient. Each dot represents the average of all mitochondria quantifications in each patient. 5 young individuals surveyed (mitochondrial number varies; Case #1: n = 253; Case #2: n = 250; Case #3: n = 250; Case #4: n = 252; Case #5: n = 253; total mitochondria surveyed across young cohort: n = 1258) and 4 old cases (mitochondrial number varies; Case #1: n = 254; Case #2: n = 250; Case #3: n = 250; Case #4: n = 250; total mitochondria surveyed across old cohort: n = 1004) for 3D reconstruction. Significance was determined with the Mann–Whitney test comparing the combined number of mitochondria in young (n = 1258) and old (n = 1004) cohorts, with **** representing p ≤ 0.0001.

### Enrollment

Specific recruitment criteria varied among cohorts and were approved by relevant institutional review boards. For all individuals from Cohorts 1, 2, 4, and 5 (Supplemental Figure 1), old participants were selected based on specific criteria, including age (50+ years), ability to engage in physical exercise, and absence of chronic diseases that could interfere with exercise. As for the younger cohort, it consisted of physical education students. These participants were in the age range of 18-50 years and were enrolled to represent the “young” demographic in our study. Their optional involvement was part of their academic curriculum, focusing on physical education and sports science. These participants were generally healthy, physically active, and had no known medical conditions that could impact the study outcomes. For Cohort 3, pre-existing biopsies were utilized with patient consent. Exclusion criteria for all cohorts: individuals with any significant cancers (e.g., solid tumors, hematological malignancies, and metastatic cancers), individuals with known co-morbidities (e.g., any existing history outside of sarcopenia), pregnant individuals or those planning to become pregnant during the study period, individuals with significant cognitive impairment or psychiatric disorders that may affect their ability to provide informed consent, individuals with severe musculoskeletal injuries affecting mobility or exercise capacity, individuals reporting active substance abuse issues, individuals who report recently participating in intensive physical training programs, individuals reporting recent surgeries, and individuals currently using medications known to significantly impact muscle structure or function (e.g., corticosteroids, statins, and neuromuscular blockers. Full patient details may be found in Supplemental Files 1-4.

### Magnetic Resonance Imaging Data

Full sample characteristics are available in Supplemental File 1 and 2. The magnetic resonance imaging (MRI) was performed on the Magnetom Essenza and Sempra, 1.5T (Siemens, Inc.) where a strong magnetic field aligns the hydrogen nuclei in the body. An RF pulse disturbs this alignment, and as the nuclei return to their original state, they emit signals that are detected and processed into detailed images in order to obtain high-resolution imaging through sequences like T2-weighted imaging, which highlights various tissues and structures. The measurement was made along the largest axis in the axial plane, in the middle third of the thigh and calves (Steinmeier et al. 1998; Eck et al. 2023).

### Segmentation and Quantification of 3D SBF-SEM Images Using Amira

The protocols followed previously established methods (Vue, Neikirk, et al. 2023; Crabtree et al. 2023; Garza-Lopez et al. 2022; Vue, Garza-Lopez, et al. 2023). Human quadriceps were excised and cut into 1 mm^3^ samples, they were fixed in 2% glutaraldehyde in 0.1 M cacodylate buffer and processed using a heavy metal protocol adapted from a previously published protocol (Courson et al. 2021; Mustafi et al. 2014). Following immersion in 3% potassium ferrocyanide and 2% osmium tetroxide for 1 hour at 4°C, the tissue was treated with filtered 0.1% thiocarbohydrazide for 20 min and 2% osmium tetroxide for 30 min, and de-ionized H_2_O washes performed between each step. Tissues were incubated overnight in 1% uranyl acetate at 4°C. Next, the samples were immersed in a 0.6% lead aspartate solution for 30 min at 60°C and then dehydrated in graded acetone dilutions. The samples were embedded in fresh Epoxy TAAB 812 hard resin (Aldermaston, Berks, UK) and polymerized at 60°C for 36–48 hours. The block was sectioned for transmission electron microscopy (TEM) to identify the area of interest, trimmed to a 0.5 mm × 0.5 mm region of interest (ROI), and glued to an aluminum pin. Finally, the pin was placed into an FEI/Thermo Scientific Volumescope 2 scanning electron microscope imaging.

Following serial imaging of the samples, 3D reconstruction of SBF-SEM orthoslices was performed using previously published techniques (Garza-Lopez et al. 2022; Hinton et al. 2023; Neikirk, Vue, et al. 2023; Vue, Garza-Lopez, et al. 2023; Vue, Neikirk, et al. 2023; Crabtree et al. 2023). Briefly, using contour tracing in Amira to perform 3D reconstruction, 300−400 orthoslices and 50–100 serial sections were stacked, aligned, and visualized. An individual blinded to the experimental conditions who was familiar with organelle morphology manually segmented the structural features on sequential slices of micrograph blocks.

### Murine-derived Myotubes

As previously described (Pereira et al. 2017), satellite cells were isolated from C57Bl/6J mice, and cells were derived and plated on BD Matrigel-coated dishes. Following our previously published protocol (Stephens et al. 2023), cells were activated to differentiate into myoblasts and myotubes.

To separate myoblasts, after reaching 90% confluence, myoblasts were differentiated to myotubes in DMEM/F-12 containing 2% fetal bovine serum (FBS) and 1× insulin-transferrin-selenium. Three days after differentiation, myotubes were infected to deliver 1 µg of CRISPR/Cas9 plasmid (Santa Cruz CRISPR Plasmid) to delete *MFN1, MFN2,* or both (double knockout, DKO), which was validated by quantitative PCR (qPCR). Experiments were performed 3–7 days after infection.

### Cell Culture

After isolation, human and primary mouse myotubes were maintained in a mixture of DMEM/F-12 (Gibco: Waltham, MA, USA) containing 20% FBS (Gibco), 10 ng/ml basic fibroblast growth factor, 1% penicillin/streptomycin, 300 µl/100 ml Fungizone, 1% non-essential amino acids, and 1 mM β-mercaptoethanol. On alternate days, the medium was replaced after cells were washed with phosphate-buffered saline (PBS) to ensure all excess media is removed.

### RNA Extraction and Real-Time qPCR

Using a RNeasy kit (Qiagen Inc.), RNA was isolated and subsequently quantified through absorbance measurements at 260 nm and 280 nm using a NanoDrop 1000 spectrophotometer (NanoDrop products, Wilmington, DE, USA). Using a High Capacity cDNA Reverse Transcription Kit (Applied Biosciences, Carlsbad CA), isolated RNA (∼1 µg) was reverse transcribed and then amplified by real-time qPCR with SYBR Green (Life Technologies, Carlsbad, CA), as previously described (Boudina et al. 2007). For each experimental condition, triplicate samples (∼50 ng DNA each) were placed in a 384-well plate before undergoing thermal cycling in an ABI Prism 7900HT instrument (Applied Biosystems). The thermal cycling conditions were set as follows:

| Cycle Count | Temperature | Time |
| --- | --- | --- |
| 1 | 95°C | 10 min |
| 40 | 95°C | 15 s |
|  | 59°C | 15 s |
|  | 72°C | 30 s |
| 1 | 95°C | 15 s |
| 1 | 60°C | 15 s |
| 1 | 95°C | 15 s |

The following primers were used **(Tezze et al. 2017)**:

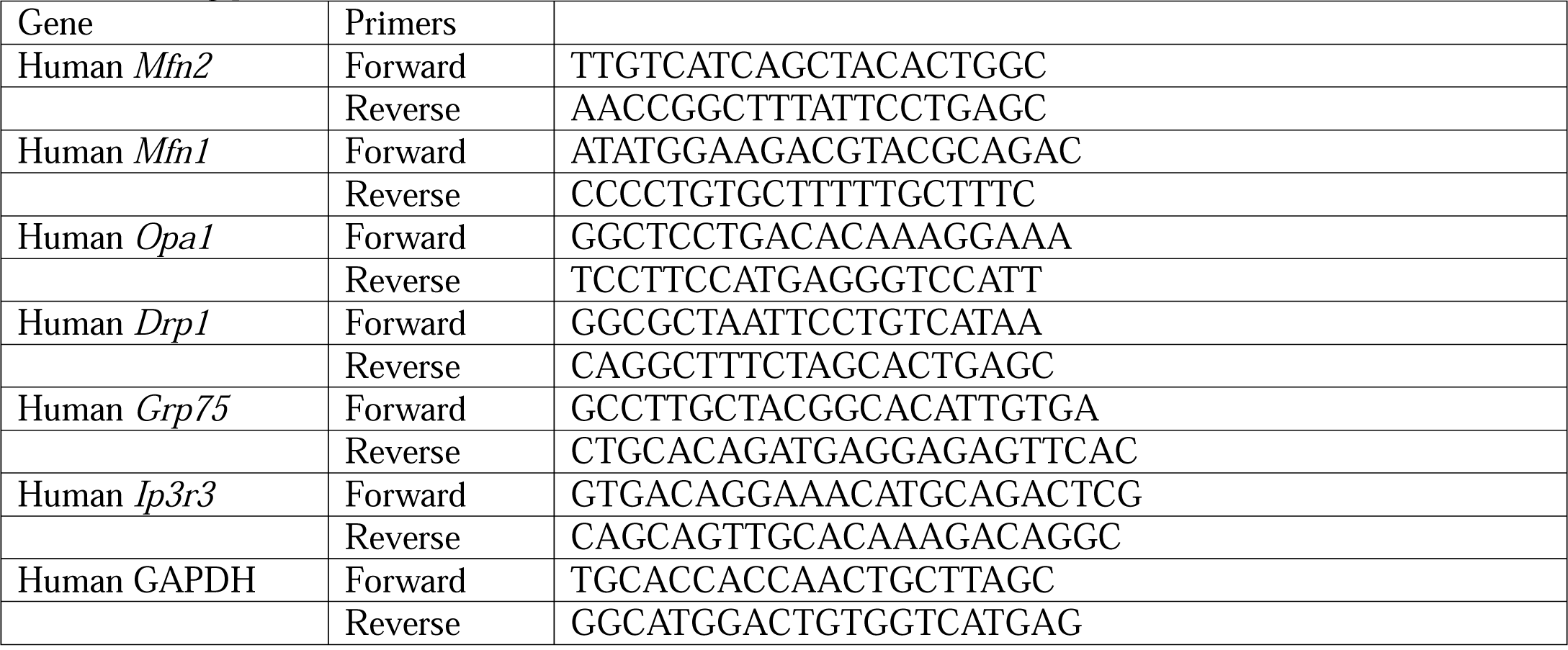

The final results are normalized to those of glyceraldehyde-3-phosphate dehydrogenase and are presented as relative mRNA fold changes.

### In Vitro Exercise Stimulation

*In vitro* exercise simulation using electric pulse stimulation was performed as previously described (Evers-van Gogh et al. 2015; Lambernd et al. 2012). Briefly, following human or skeletal myotube differentiation or with C2C12 cells, the cells were starved by culturing in DMEM without FBS. The medium was replaced directly prior to stimulation. Electrical stimulation was administered through carbon electrodes using a C-Pace 100 pulse generator (IonOptix, Milton, MA, USA) in a C-dish. The stimulation parameters were set at a frequency of 1 Hz, a pulse duration of 2 ms, and an intensity of 11.5 V, with treatment sustained for 4.5 or 24 hours. Conditioned medium was harvested from both stimulated and non-stimulated conditions and then centrifuged at 800 rpm/17 rcf for 5 min, and the samples were stored at −80°C. A mixture of growth medium DMEM/F-12 containing 10% FBS and conditioned medium containing 0% FBS in equal parts was prepared and applied to specified cell lines. Western blotting or qPCR was then performed as described.

### Western Blotting

As previously described (Hinton et al. 2024), to obtain protein extracts from differentiated myotubes and C2C12 cells, we washed cells with ice-cold PBS and then added cold lysis buffer [25 mM Tris HCl, pHL7.9, 5LmM MgCl_2_, 10% glycerol, 100 mM KCl, 1% NP40, 0.3LmM dithiothreitol, 5 mM sodium pyrophosphate, 1 mM sodium orthovanadate, 50LmM sodium fluoride, and protease inhibitor cocktail (Roche Applied Science, Penzberg, Germany)]. Following scraping of the cells, a 25-gauge needle was used to homogenize the cells before they were centrifuged at 14,000 rpm/5,268 rcf for 10 min at 4°C. Following centrifugation, the supernatants were collected and diluted with Laemmli sample buffer to obtain a final concentration of 1×. We performed sodium dodecyl sulfate-polyacrylamide gel electrophoresis with 1× concentrated cell lysates, and proteins were transferred to nitrocellulose membranes (BioRad, Berkeley, California, USA). Blocking of membranes was performed with 5% bovine serum albumin in Tris-buffered saline with Tween-20. Primary antibodies used for western blotting and their working dilutions included calreticulin (CALR) and fibroblast growth factor 21 (FGF21) (1:1000 diltuion, Abcam). Following incubation of three biological replicates for each protein of interest, quantification was performed using Image Studio Lite Ver 5.2.

### Transmission Electron Microscopy (TEM) Analysis

As previously described (Hinton et al. 2023), cells were fixed in 2.5% glutaraldehyde diluted in sodium cacodylate buffer for 1 hour at 37°C and then embedded in 2% agarose, postfixed in buffered 1% osmium tetroxide, stained with 2% uranyl acetate, and dehydrated with a graded ethanol series. Following EMbed-812 resin embedding, 80-nm sections were cut on an ultramicrotome and stained with 2% uranyl acetate and lead citrate. Images were acquired on a JEOL JEM-1230 transmission electron microscope operating at 120 kV.

NIH ImageJ software (Schneider et al. 2012) was used to manually trace and analyze all mitochondria or cristae using the freehand tool (Parra et al. 2013). Measurements of mitochondrial area, circularity, and number were performed using the Multi-Measure ROI tool in ImageJ (Lam et al. 2021; Neikirk, Vue, et al. 2023; Parra et al. 2013). We used three distinct ROIs, all of the same magnification, in ImageJ to examine cristae morphology and determine their area and number. The sum of the total cristae area divided by the total mitochondrial area was used as a proxy to determine cristae volume (Patra et al. 2016).

### Mitochondrial Area and Circularity Analysis

Mitochondrial morphology was assessed by quantifying mitochondrial circularity and area, which were measured using ImageJ software. The mitochondrial area was measured for every mitochondrion in the region. Circularity is a measure of how closely a shape approximates a perfect circle, calculated as 4π × (area/perimeter^2^). A value of 1.0 indicates a perfect circle. Increased mitochondrial circularity indicates a shift toward more rounded mitochondria and loss of elongated mitochondrial networks. Graphs were created and statistical analysis was performed using GraphPad Prism (version 9.0, La Jolla, CA, USA).

### Drosophila Strains and Genetics

Flies were cultured on standard yeast-cornmeal agar medium in vials or bottles at 25°C with a 12-hour light/dark cycle. The *Mef2*-*Gal4* (also known as P{GAL4-Mef2.R}3) driver line was used to direct the expression of upstream activating sequence (UAS) transgenes, specifically in skeletal muscle. UAS-mitoGFP (II) was used to visualize mitochondria. RNAi knockdown (KD) lines originating from transgenic RNAi lines were obtained from the Bloomington *Drosophila* Stock Center and included *UAS-Marf RNAi* (55189). Chromosome designations and additional strain details are available on FlyBase (http://flybase.org). Male and female flies were analyzed together, as no sex differences in mitochondrial morphology were observed in wild-type muscle. The Mef2-Gal4 strain served as a control within the respective genetic backgrounds.

### Mitochondrial Staining

Thoraces from 2–3-day-old adult *Drosophila* were dissected in 4% paraformaldehyde (PF, Sigma), and indirect flight muscles were isolated as described previously (Katti et al. 2022). Isolated muscles were fixed in 4% PF for 1.5 hours with agitation, followed by three 15-min washes with PBSTx (phosphate-buffered saline + 0.3% Triton X-100). Mitochondria were visualized by staining with either 200 nM for MitoTracker™ Red FM (M22425, ThermoFisher) for 30 min or by mitochondrial-targeted green fluorescent protein (GFP) expressed from UAS-mito-GFP under the control of Mef2-Gal4. F-actin was stained by incubating muscles in 2.5 μg/ml phalloidin-TRITC (Sigma) in PBS for 40 min at 25°C.-Stained muscles were mounted in Prolong Glass Antifade Mountant with NucBlue (ThermoFisher) and imaged using a Zeiss LSM 780 confocal microscope.

### Mitochondrial Quantification

Mitochondria were quantified by imaging muscle fibers using fluorescence microscopy. MitoTracker Green FM dye (Invitrogen) or mito-GFP, as mentioned above, was used to label mitochondria. Images were acquired at 60× magnification and were analyzed using ImageJ software. Images were divided into regions, and mitochondria spanning three sarcomeres (from the Z-disc of the first sarcomere to the Z-disc of the fourth sarcomere) were selected for analysis. The number of mitochondria in three sarcomeres was manually counted using ImageJ.

### RNA Sequencing

Using the same method of RNA isolation as described above, for RNA sequencing, a list differentially expressed genes (DEGs) was compiled from RNA-sequencing results (p_adj_<0.05 and absolute log_2_ fold change>0.66) and were analyzed for potential enriched pathways using Ingenuity Pathway Analysis (IPA, QIAGEN) and Gene Set Enrichment Analysis (GSEA) with WebGestalt (www.webgestalt.org) (Liao et al. 2019). For IPA analysis, enriched pathways were considered significant when applying an absolute activation Z-score of >2 and p_adj_<0.05. For GSEA results, an absolute enrichment score of >2 and p_adj_<0.05 was considered significant.

### Data Analysis

Black bars in graphs represent the standard error, and dots represent individual data points. All analyses were performed using the GraphPad Prism software package, with specific tests indicated in the figure legends. A minimum threshold of p < 0.05 indicated a significant difference (as denoted by *). Higher degrees of statistical significance (**, ***, ****) are defined as p < 0.01, p < 0.001, and p < 0.0001, respectively.

## Results

### Human Aging Causes Alterations in Muscle Size

Previous studies have utilized magnetic resonance imaging of thigh cross-sectional area (CSA) as a proxy to determine muscle size (Beneke et al. 1991). Furthermore, the muscle and bone relationship has been examined to investigate sarcopenia because the intra-individual muscle mass loss during aging can be determined (Maden-Wilkinson et al. 2014). Thus, we initially utilized magnetic resonance imaging to determine how the skeletal muscle structure in the thigh and femur is remodeled during the aging process. By enrolling female and male participants (Figures 1A–D), we created a “young” cohort consisting of individuals from 18 to 50 years old and an “old” cohort of individuals older than 50 years old (Supplemental File 1). When male and female participants were combined, thigh or femur CSA was not significantly differentiated across the aging process (Supplemental Figure 1A). However, we observed slight sex-dependent differences during the aging process (Supplemental Figures 2B–C). Males had significantly decreased thigh CSA (Figure 1E-F), while females had increased femur CSA (Figures 1G–H). For both sexes, however, the muscle area relative to the bone area in the thigh region generally decreased (Figures 1I–J). We proceeded to look at calf measurements in a new cohort of individuals (Supplemental File 2) (Figures 1A’-D’). Looking at metrics including the tibia and total calf CSA, we observed no sex-dependent differences during the aging process (Supplemental Figures 2D–E). Tibia CSA, total muscle CSA, and the ratio of these measurements were statistically unchanged except for females showing a slight age-dependent loss in total calf muscle CSA (Figures 1E’-J’). Together, these age-related losses in muscle mass demonstrated the occurrence of muscle atrophy in the human quadriceps. While we could not confirm participants had sarcopenia, these results support the observation of an age-related decline in muscle mass. Next, we sought to explicate whether these age-related losses in muscle mass are correlative with alterations in 3D mitochondrial structure.

### Aging is Associated with Changes in Human Skeletal Muscle Mitochondrial Structure and Dynamics

Based on the age-related loss of thigh CSA, we sought to understand how the aging process may lead to this degeneration of muscle mass. Since mitochondrial size and morphology are dynamic, responding to environmental stimuli and conferring changes in the metabolic effects of mitochondria such as respiratory efficiency (Frey & Mannella 2000; Glancy et al. 2020), we first sought to determine whether mitochondrial dynamics and morphology are altered as a result of the aging process. Utilizing qPCR, we surveyed pertinent mitochondrial proteins from skeletal muscle samples of 18–50-year-old (young) and 50–90-year-old (old) humans. In human skeletal muscle, genes associated with mitochondrial fusion and fission dynamics (Figures 2A–D) and MERCs (Figures 2E–G) were altered with aging. Specifically, fusion (Mfn1, Mfn2, and Opa1; Figures 2A–C) and fission (Drp1; Figures 2D) proteins were decreased with aging. Reductions in fusion and fission proteins were confirmed by previous studies that showed age-related losses of mitochondrial dynamic proteins in human skeletal muscle (Seo et al. 2010; Crane et al. 2010). Interestingly, however, MERC proteins (Grp75; Ip3r3, and Vdac3; Figures 2E–G) were upregulated in human skeletal muscle during aging. These results are suggestive of decreases in mitochondrial dynamics and mitochondrial structural integrity; however, the exact 3D structural remodeling of mitochondria in human skeletal muscle remains unclear. Therefore, we sought to understand how these changes in mRNA transcripts manifested as alterations in mitochondrial structure.

We utilized SBF-SEM to perform 3D reconstruction of mitochondria (Figure 2H). SBF-SEM has a lower resolution than conventional TEM imaging (Marshall, Neikirk, et al. 2023; Neikirk, Lopez, et al. 2023), but its high range offers advantages over other 3D light-based imaging methods (Marshall, Damo, et al. 2023; Marshall, Krystofiak, et al. 2023). We collected quadriceps samples from young (under 50 years old) and old (over 50 years old) individuals (Supplemental File 3). We utilized a mix of human quadriceps from both males and females because previous studies have generally shown that sarcopenia occurs at a similar rate in both sexes, and we observed only a few significant sex-dependent differences in age-dependent loss of muscle (Tay et al. 2015). To begin, we qualitatively looked at MERCs in our samples. While there is some controversy, many studies demonstrate MFN2 also acts as a MERC tether protein in addition to its role as a mitochondrial dynamic protein (Han et al. 2021; Sebastián et al. 2012; Basso et al. 2018). Since our data also showed the upregulation of MERC proteins, we wanted to briefly understand how MERCs changed. Qualitatively, when comparing young (Figure 2I) to old (Figure 2J), we saw that with aging MERCs appeared smaller contact area, but also interacted with more mitochondria (Figure 2K). However, we also noticed differences in mitochondrial phenotypes, so we decided to perform a rigorous analysis of mitochondria.

According to previous methods (Garza-Lopez et al. 2022), 50 z-directional SBF-SEM orthogonal micrographs, also known as “orthoslices”, were obtained (Figure 2L–M), and the 3D structure of intermyofibrillar mitochondria, or mitochondria between fibrils (Vendelin et al. 2005), was rendered through manual contour segmentation (Figure 2L’–M’). This time-consuming manual process allowed for mitochondria structure to be verified and for observation of the complete mitochondrial 3D structure in young and old human skeletal muscle (Figure 2L’’-M’’). Five ROIs were considered in the young condition and 4 ROIs were considered in the old condition, within which approximately ∼250 mitochondria were quantified for a total of ∼2250 mitochondria (Supplemental Figure 3).

Once rendered, we found no significant change in the mitochondrial surface area or perimeter, but interestingly the mitochondrial volume increased in aged samples (Figure 2O–P). Surface area and perimeter unchanging refers to overall stability in mitochondrial outer dimensions. However, mitochondrial volume can be indicative of the total capacity for energy production (Gallo et al. 1982), it can also be indicative of mitochondrial swelling, an event that typically occurs antecedent to apoptosis (Safiulina et al. 2006). Thus, to better understand how mitochondrial structure undergoes age-related changes, we also examined mitochondrial complexity.

### Aging Causes Human Skeletal Muscle Mitochondria to Become Less Complex

Mitochondrial complexity has been shown to be altered with mtDNA defects in human skeletal muscle (Vincent et al. 2019), yet how the 3D complexity changes with aging remains unclear. Based on structural changes in surface area, we sought to determine whether aging conferred a modulatory effect on mitochondrial complexity. Using the same samples as before, intermyofibrillar mitochondria from young and old human participants were viewed from transverse (Figures 3A–B) and longitudinal points of view (Figures 3A’–B’). Based on the views of mitochondria from these disparate axes, we observed that mitochondria from young individuals generally appeared more elongated, while those from older individuals appeared more compact. To confirm this finding, we examined mitochondrial sphericity, which showed age-related increases, in which mitochondria generally appeared to have a rounder shape (Figure 3C). As a secondary 3D form-factor measurement, we employed a mitochondrial complexity index (MCI), which represents the ratio of the surface area and volume (Vue, Garza-Lopez, et al. 2023; Vincent et al. 2019). The MCI indicated reduced complexity in samples from older individuals (Figure 3D). To further visualize how these reductions in complexity arise, we used Mito-otyping, a method of mitochondrial organization based on their relative volume, to visualize changes in the complexity (Vincent et al. 2019). Mito-otyping showed that, in young individuals, mitochondria were more complex and showed diverse phenotypes, while older individuals had mostly compact and spherical mitochondria (Figure 3E). These changes also indicate that the surface area to volume ratio decreases during the aging process, since aged samples have a higher volume without any change in surface area. While slight intra-cohort variability in the MCI was observed (Supplemental Figure 3F), the relatively low intra-individual variability indicated that changes in mitochondrial complexity and morphology were generally ubiquitous across the mitochondria surveyed. Notably, Mito-otyping also allows for characteristics of the sample population to be compared (Supplemental Figure 3A; Videos 1-9), but we generally found no hallmarks associated with specific sexes. For example, the two females in the young cohort (Supplemental File 3; Supplemental Figure 3; Young Case #1 and Young Case #2) display very different overall phenotypes: one marked by highly complex mitochondria and another marked by spherical mitochondria (see the top 2 rows of Mito-otyping). Similarly, in the old cohort, the male surveyed (Old Case #3) presents a similar phenotype to two of the other females (Old Case #1 and Old Case #2), suggestive of interindividual heterogeneity owing to inter-cohort differences more so than sex-dependent differences. Together these results indicate that mitochondrial volume is significantly increased, potentially as a sign of swelling in aged human vastus lateralis, thigh, and quadriceps, while the mitochondrial morphology undergoes significant alterations, which may be representative of mitochondrial dysfunction and associated sarcopenia.

### Aging Modulates Exercise Ability, Immune and Glucose Responses

Next, we sought to understand the functional implications of these age-related changes. Because no pharmacological interventions for sarcopenia yet exist, exercise has remained a principal therapeutic approach to mitigate this condition (Phu et al. 2015), and generally can be an important mechanism against other age-related muscle weaknesses. Inversely, however, the muscle mass loss caused by sarcopenia can reduce endurance and strength, which suggests that adequate exercise before aging may be necessary (Greenlund & Nair 2003). We sought to understand how individuals may have impaired strength and endurance. We included a new cohort of individuals constituting both males and females, divided them into “young” (under 50 years old) or “old” (over 50 years old) categories, and subjected them to various exercises (Supplemental File 4). A walking test (Supplemental Figure 4A), a grip strength test (Supplemental Figure 4B), a test of localized muscle endurance (LME) of the lower body per modified protocols (Jones et al. 1999) (Supplemental Figure 4C; Video 10), and a test of LME of the upper body through an adapted method (Sato et al. 2021) (Supplemental Figure 4D; Video 11) were performed. We observed lower exercise strength and endurance in older individuals with slight sex-dependent differences (Supplemental Figure 5). Despite this decline, there were minimal changes in weight and body mass index when comparing young and old individuals. This suggests that changes in strength are not primarily attributable to alterations in overall mass (Figures 4A–B’). Both males and females also had significant decreases in walking distance and the associated maximum amount of oxygen utilized during intense exercise (VO_2_) suggesting decreased aerobic capacity (Figures 4C–D). Notably, this difference was slightly more pronounced in females than males (Figures 4C’–D’). When examining strength, we showed that the grip strength of both arms was lower in aged samples, indicating decreased muscle mass (Figures 4E–F’). Interestingly, while young males had greater grip strength than females, males also exhibited a more significant decrease with aging, resulting in aged males and females having similar grip strength (Supplemental Figures 5F–G). To further explore how muscle endurance changes, we examined both upper and lower body endurance, which showed much more drastic decreases in lower body endurance, with slight sex-dependent differences (Figures 4G–H’). Together, these results indicate that with aging, although a distinct cohort of individuals, both males and females lose endurance and muscle strength, potentially indicative of age-related atrophy occurring correlatively with the 3D structural mitochondrial remodeling that we observed.

**Figure 4.**
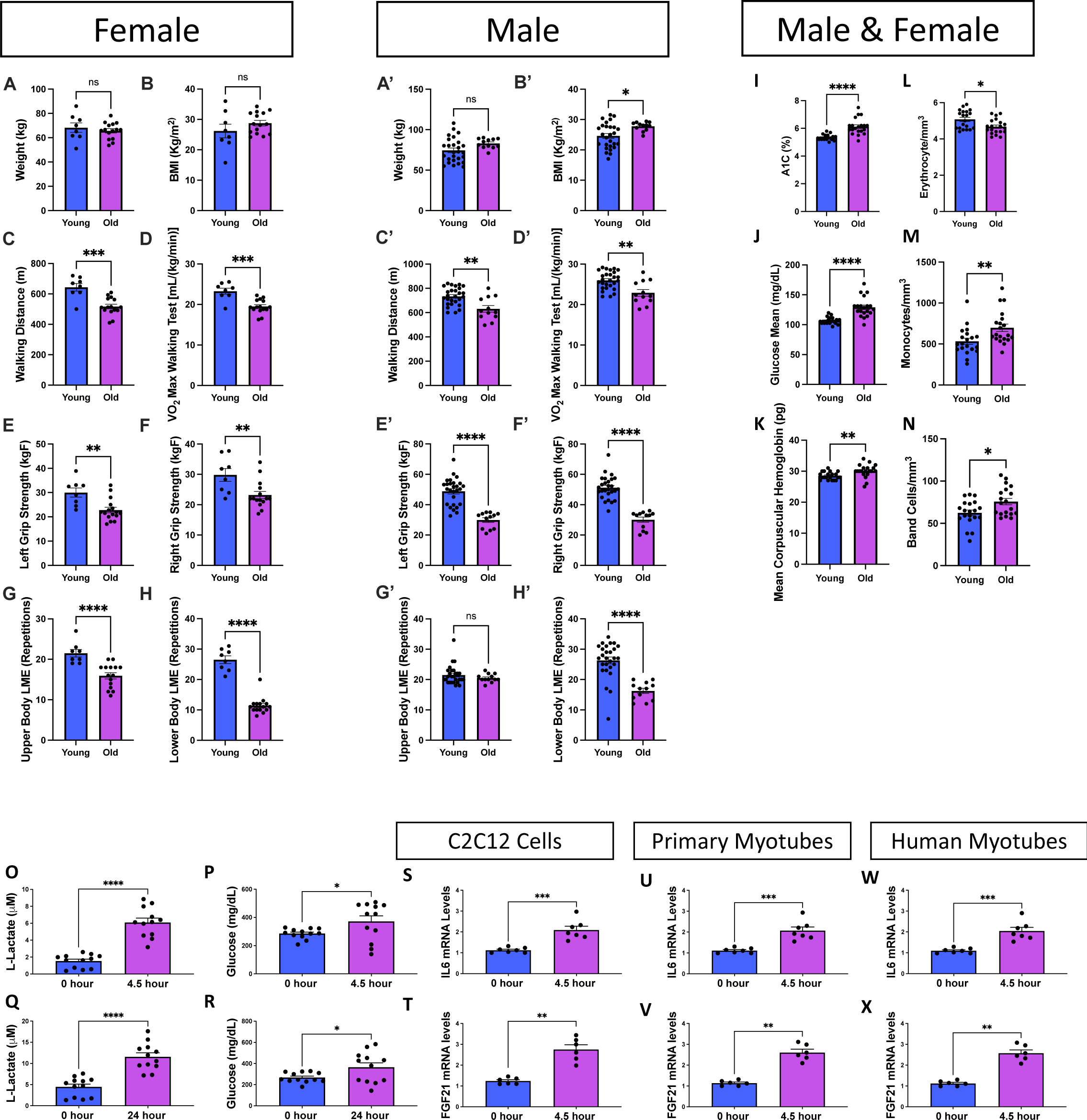
Aging Changes Exercise Parameters Associated with Immune Modulatory Functions. Exercise data from (A–H) females under 50 years old (aged 21–26 years; n = 8), females over 50 years old (aged 60–73 years; n = 15), (A’–H’) males under 50 years old (aged 19–35 years; n = 27), and males over 50 years old (aged 63–76 years; n = 12). Blue bars represent young individuals, and purple bars represent older individuals. (A–B) Plots detailing weight and body mass index distribution of females and (A’–B’) of young and old individuals, (C–C’) walking distances (in meters), and (D–D’) VO_2_ max values during a walking test among the same groups. (E–F) Scatter box plots for grip strength in kg. (E) Left grip strength and (F) right grip strength for females and (E’–F’) males. (G–H) Plots representing localized muscle endurance (G) of the lower body and (H) the upper body across females and (G’–H’) males. (I– N) Molecular and physiological measurements from the plasma of male and female participants; the full analysis is shown in Supplemental Figure 4. (I) Glycated hemoglobin (A1C) percentage levels in young and old participants. (J) Glucose concentration levels, presented in milligrams per deciliter (mg/dl), in young and old participants. (K) Average concentration of hemoglobin in a given volume of packed red blood cells, known as the mean corpuscular hemoglobin (MCH), measured in picograms (pg) for both age groups. (L) Erythrocyte (red blood cell) count measured in millions per cubic millimeter (mm^3^) for young and old participants. (M) Monocyte count, depicted as cells per cubic millimeter (cells/mm^3^), for young and old participants. (N) Band cell (immature white blood cell) count measured in cells/mm^3^ for young and old participants. (O) *In vitro* exercise stimulation in human myotubes with L-lactate and (P) glucose quantification after 4.5 and (Q–R) 24 hours. IL6 mRNA levels, as determined by quantitative PCR, are shown for (S) C2C12 cells, (U) primary myotubes, and (W) human myotubes. FGF21 mRNA levels, as determined by quantitative PCR, are shown for (T) C2C12 cells, (V) primary myotubes, and (X) human myotubes. Each dot represents an individual patient (Supplemental File 2) or experimental run. Significance was determined with the Mann–Whitney test, with **** representing p ≤ 0.0001.

To further understand potential mechanisms underpinning age-related changes, we determined whether any factors in blood or plasma exhibited alterations. Within a new cohort (Supplemental File 5); We found that aging had numerous effects on blood serum molecules (Supplemental Figure 6). Of these, changes in glucose metabolism, hemoglobin-carrying capacity, and immune responses were notable (Figures 4I–N). Glycated hemoglobin levels, which increased with aging (Figure 4I), are used to gauge average blood glucose over approximately the previous 3 months and serve as a better index for long-term glycemic exposure than fasting or blood glucose levels (Zhang et al. 2010). Similarly, in the older cohort, the mean glucose concentration was significantly increased, suggesting impaired glucose homeostasis with age (Figure 4J). When investigating the mean corpuscular hemoglobin value, which signifies the average amount of hemoglobin in red blood cells, we noticed a significant increase in older individuals (Figure 4K). This finding is suggestive of increased oxygen-carrying capacity, which can affect muscle endurance and function and may be a response to altered mitochondrial respiration (Xuefei et al. 2021). While alterations in hemoglobin can affect mitochondria through oxidative stress generation (Anon 2021), hemoglobin serves multifaceted functions, and peripheral blood mononuclear cells can also increase intracellular hemoglobin (Brunyanszki et al. 2015). Thus, we next investigated how aging increases the immune response. As expected, because autoantibody responses generally increase in the elderly population (Yung 2000), we found that while erythrocytes were decreased, monocytes and band cells were both increased in the aged group compared with those in the young group. Notably, an elevated band cell count suggests an active inflammatory or infectious process (Mare et al. 2015). Together, these findings show that glucose levels increase concomitantly with immune responses during the human aging process.

Next, we sought to determine whether exercise, which has extensively been described as an effective therapy for sarcopenia (Phu et al. 2015; Taaffe 2020), mechanistically acts to modulate these age-dependent alterations in immune signaling and glucose metabolism. To explore this mechanism, we used a previously established method of electric pulse stimulation, which simulates exercise *in vitro* (Evers-van Gogh et al. 2015). We found that with *in vitro* electrical stimulation for 4.5 hours, L-lactate and glucose levels both increased in human myotubes (Figures 4O–P), confirming that cells exhibit an “exercised” phenotype. Specifically, lactate can serve as a valuable energy source, but its accumulation can also be linked to muscle fatigue (Nalbandian & Takeda 2016). Increased glucose levels reflect the body’s reliance on carbohydrates for energy during exercise (Mul et al. 2015). We further recapitulated these findings with 24 hours of electrical stimulation, which showed no significant differences and suggested that 4.5 hours is sufficient to create an exercised phenotype (Figures 4Q–R). Once we confirmed this method of *in vitro* exercise, we examined three cell types: C2C12 cells, primary myotubes, and human myotubes. In all three cell types, interleukin 6 (IL6; representative of the immune and inflammatory response) and FGF21 (representative of glucose metabolism) both increased with exercise (Figures 4S–X). Notably, this increase in IL6 is consistent with previous studies (Beavers et al. 2010), suggesting an acute response that may not be chronic. However, this increase does not suggest that exercise mitigates sarcopenia through reductions in the immune response. Notably, however, FGF21 is well understood to be antihyperglycemic, with increases in circulating FGF21 levels occurring concomitantly with improved glucose tolerance and decreased blood glucose (Xu et al. 2009). Beyond broad associations of mitochondrial dysfunction with insulin sensitivity, FGF21 has previously been implicated in interactions with mitochondrial dynamic proteins (Pereira et al. 2017; Pereira et al. 2021). Together, these findings demonstrate that exercise has a modulatory effect on age-related changes in glucose homeostasis, indicating that aging effects can be modulated in part through exercise. Notably, this finding is suggestive that exercise mitigates age-related muscle atrophy, with implications for potential FGF21-dependent pathways, which may involve mitochondrial dynamic alterations in aging and sarcopenia. Thus, we turned our attention to understanding whether mitochondrial structure can be modulated by exercise.

### Exercise Restores Age-related Loss of Mitofusin 2

As previously shown, exercise may be an important mechanism by which mitochondrial quality is protected as healthy aging, including the mitigation of sarcopenia, occurs (Cartee et al. 2016). In particular, lifelong endurance exercise training enhances mitochondrial volume, network connectivity, and oxidative capacity in older human skeletal muscle, as compared to untrained and moderately trained older humans (Ringholm et al. 2023). It has also previously been shown that PGC-1α, a key factor of mitochondrial biogenesis, is increased following exercise (Koh et al. 2017; Wright et al. 2007; Baar et al. 2002), but we sought to better explicate changes in regulators of mitochondrial dynamics.

Past seminal findings have shown that MFN2 plays a critical role in muscular aging and associated mitochondrial dysfunction through age-dependent loss of MFN2 and concomitant inhibition of mitophagy (Sebastián et al. 2012). We sought to extend these findings to determine whether MFN2 further plays a critical role in mitochondrial structure remodeling in exercise to protect against sarcopenia. Consistent with previous findings, we found that loss of MFN2 occurs in various murine tissue types during aging (Figures 5A–D), which parallels the loss we observed previously in human skeletal muscle (Figure 2B). Thus, we aimed to determine whether exercise can rescue these levels (Figures 5E–E’), which would suggest a reversal of age-related deficits in mitochondrial dynamics and structure. Again, subjecting both C2C12 (Figure 5F–I) and primary murine-derived myotubes (Figures 5F’–I’) to *in vitro* electric stimulation, we verified the ability of our *in vitro* method to induce a prolonged exercise phenotype. We observed an increase in L-lactate levels and a decrease in glucose levels following 4.5 or 24 hours of exercise (Figures 5F–I’), which may reflect a metabolic state associated with high-intensity or prolonged exercise. Specifically, this finding suggests a shift toward anaerobic metabolism, muscle fatigue due to acidosis, and the utilization of glucose and glycogen for energy (Goodwin et al. 2007). Western blotting (Figures 5E–E’) also indicated that MFN2 and CALR levels are increased in both C2C12 and primary myotubes (Figures 5J–K’), indicating that mitochondrial and MERC dynamics are increased following exercise. MFN2 and CALR are interestingly both MERC proteins (Han et al. 2021; Peggion et al. 2021), but MFN2 also plays a pluralistic role in mitochondrial fusion.

**Figure 5.**
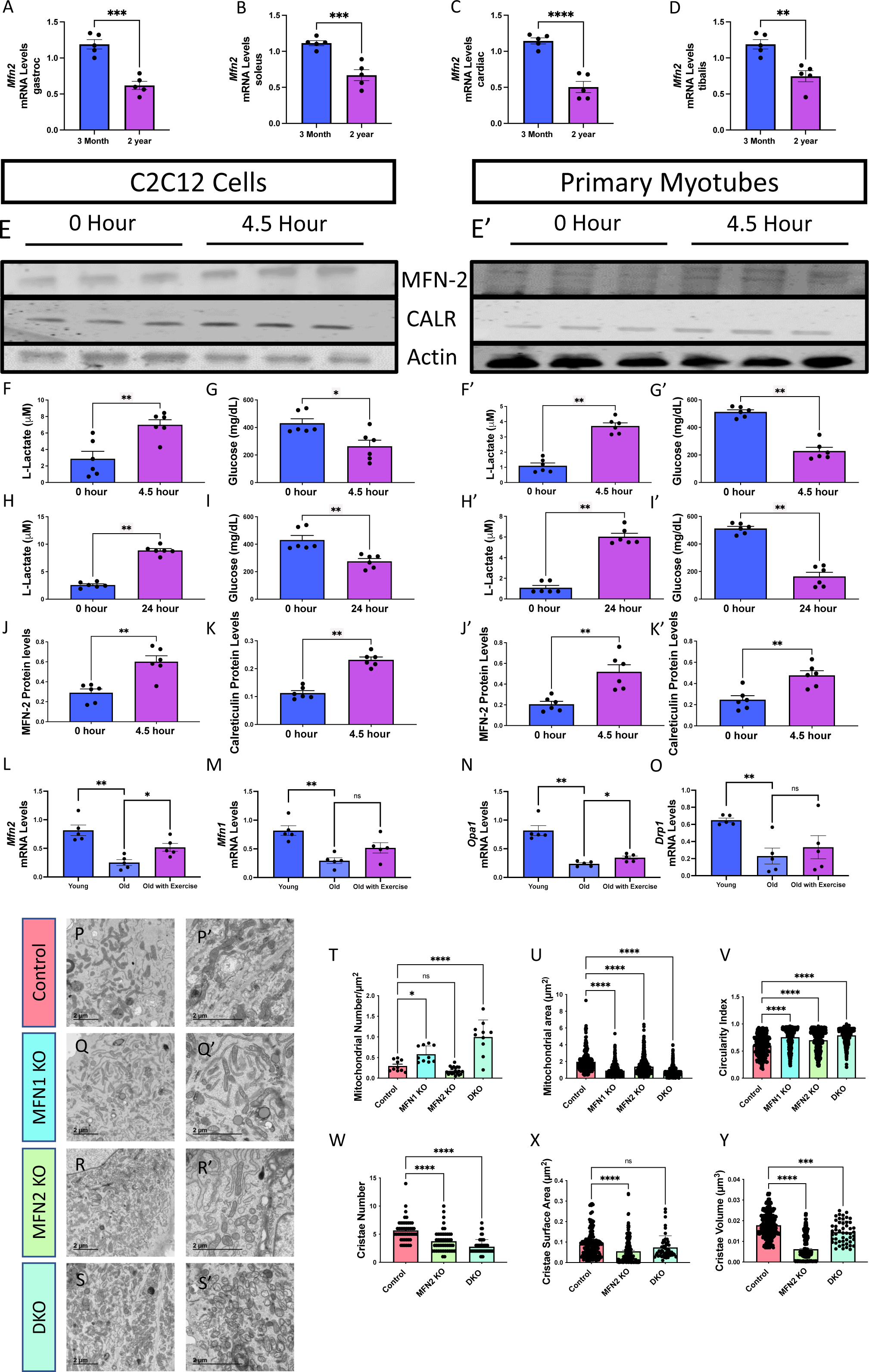
Mitofusin 2 (MFN2) Expression Changes in Response to Exercise and Aging and Changes in Mitochondrial Morphology. (A) Bar graphs show the mRNA levels (n=5), as determined by quantitative PCR, of *Mfn2* at two distinct time points, 3 months and 2 years, from murine soleus tissue, (B) gastrocnemius tissue, (C) the tibialis, and (D) cardiac tissue. (E) Western blot analysis of MFN2, calreticulin (CALR), and actin protein levels in C2C12 cells and (E’) primary myotubes after *in vitro* exercise stimulation at two-time intervals, 0 and 4.5 hours. (F–G) Quantitative analysis of lactate and glucose levels in C2C12 cells and (F–G’) primary myotubes after *in vitro* exercise stimulation for 4.5 hours. (H–I) Quantitative analysis of lactate and glucose levels in C2C12 cells and (H–I’) primary myotubes after *in vitro* exercise stimulation for 24 hours. (J) Quantification of MFN2 protein levels, normalized to actin levels, after 4.5 hours of *in vitro* exercise stimulation in C2C12 cells and (J’) primary myotubes. (K) Quantification of CALR protein levels, normalized to actin levels, after 4.5 hours of *in vitro* exercise stimulation in C2C12 cells and (K’) primary myotubes. (L) Bar graphs show the mRNA levels (n = 5), as determined by quantitative PCR, of *Mfn2,* (M) *Mfn1*, (N) *Opa1*, and (O) in *Drp1* mRNA transcripts in a distinct group of young humans (under 50 years old), old humans who do not report regular exercise of 2-3 sessions per week (over 50 years old), and old humans who regularly report life-long regular exercise of 2-3 sessions per week (over 50 years old). (P– S’) Transmission electron microscopy (TEM) images from murine-derived skeletal muscle myotubes highlighting mitochondrial morphology under different conditions: (P–P’) control, (Q-Q’) Mitofusin 1 knockout (MFN1 KO), (R–R’) Mitofusin 2 knockout (MFN2 KO), and (S–S’) double knockout (DKO). (T–U) Quantitative representation of (T) mitochondrial number average per cell, (U) mitochondrial area, and (V) mitochondrial circularity in cells under control, MFN1 KO, and MFN2 KO conditions. (W–Y) Quantitative representation of (W) cristae number, (X) cristae volume, and (Y) cristae surface area in cells under control, MFN1 KO, and MFN2 KO conditions. Each dot represents an individual mitochondrion for TEM data with variable sample number [mitochondrial number: n = ∼10; mitochondrial area: n = 296 (Control), 583 (MFN1 KO), 466 (MFN2 KO), and 999 (DKO); circularity index: n = 296 (Control), 583 (MFN1 KO), 466 (MFN2 KO), and 999 (DKO); cristae score: n = ∼50; cristae surface area: n = 192 (control), 192 (MFN2 KO), and 50 (DKO); cristae volume: n = 432 (control), 432 (MFN2 KO), and 50 (DKO)]. Intergroup comparisons were performed using either (A-K’) Mann– Whitney test or (L-Y) one-way ANOVA with Dunnett’s multiple comparisons test *post hoc*. Statistical significance is denoted as ns (not significant), *p < 0.05, **p < 0.01, ***p < 0.001, or ****p < 0.0001.

To recapitulate these findings in a human model, we looked at mRNA transcript levels of fusion and fission proteins *Mfn2, Mfn1*, *Opa1*, and *Drp1*, in three distinct groups: young humans (under 50 years old), old humans who do not report regular exercise of 2-3 sessions per week (over 50 years old), and old humans who regularly report life-long regular exercise of 2-3 sessions per week (over 50 years old) (Figures 5L-O). Concurrently with previous studies (Sharma et al. 2019; Srivastava 2017), all of these mitochondrial dynamic proteins decreased with aging; yet, the old cohort reporting exercise only shows significant increases in *Mfn2* and *Opa1* mRNA transcripts.

Because we noted that exercise may reverse age-related loss of MFN2, next we focused on elucidating the role of MFN2 in mitochondrial structure. While past studies have consistently shown that MFN2 deficiency is associated with increased MERC tethering (Leal et al. 2016; De Brito & Scorrano 2008), some studies have shown that MFN2 modulation does not necessarily impact mitochondrial structure alone (Cosson et al. 2012). In some species, such as *Drosophila melanogaster*, a single protein is functionally analogous to both MFN1 and MFN2 (Dorn et al. 2011; Katti et al. 2021). Thus, to better understand the structural impacts of age-related mitofusin loss, we knocked out both MFN1 and MFN2 individually as well as contiguously (DKO) and utilized TEM analysis (Lam et al. 2021) to consider ultrastructural changes in mitochondria and cristae in murine-derived myotubes (Figures 5P–S’). MFN1 knockout (KO) and DKO resulted in increased mitochondrial numbers and decreased average mitochondrial area (Figures 5P–U). Similarly, MFN2 KO reduced mitochondrial area but inversely increased mitochondrial number, resulting in no significant change in mitochondrial count, which trended downwards (Figures 5T–U). Notably, all conditions caused the circularity index to increase (Figure 5V), resulting in more regularly shaped small mitochondria, suggestive of reduced fusion. Loss of fusion proteins may also affect cristae ultrastructure (Vue, Neikirk, et al. 2023); therefore, we specifically investigated MFN2 and the DKO condition as regulators of cristae morphology. MFN2 KO resulted in reductions in cristae number, volume, and surface area, suggesting a reduced oxidative capacity (Figured 5W–Y). The same effects on mitochondrial and cristae morphology were observed in the DKO condition, marked by smaller, more plentiful mitochondria with reductions in cristae count and volume (Figure 5W–Y).

These structural changes confirmed prior studies indicating that the loss of MFN1 and MFN2 results in structural alterations with redundant yet distinct roles (Chen et al. 2003). The results wer also consistent with previous literature showing that age-related losses of MFN1 and MFN2, along with other proteins associated with biogenesis, are reversed by exercise training (Koltai et al. 2012). Furthermore, the age-related loss of both MFN1 and MFN2 (Figures 2A–B) suggests that structural rearrangements in mitochondria may arise due to the loss of MFN1 and MFN2. Additionally, exercise may be able to ameliorate some of the age-related structural losses in mitochondria through increasing MFN2 (Figures 5J–J’; Figures 5L). However, to establish the metabolic impacts of mitofusins and determine whether the structural rearrangements caused by mitofusins are evolutionarily conserved, we examined a *Drosophila* model.

### Mitofusins are Functionally Required for Mitochondrial Regulation and Structure

We focused on flight skeletal muscle in the *Drosophila* model (Figure 6A). We knocked down Mitochondrial Assembly Regulatory Factor (Marf KD), which is functionally analogous to MFN1 and MFN2, and verified that the gene was silenced at the mRNA transcript level (Figure 6B). Then, we showed that the loss of Marf altered overall development (Figure 6C), motor skills (Videos 12-15), and fly steps, or walking motion, (Figure 6D) in *Drosophila*. Furthermore, RNA sequencing was performed in Marf KD muscle that revealed widespread transcriptional changes including Marf (Figure 6E). Pathway analysis for contributing canonical pathways using IPA indicates a primary defect in mitochondrial and metabolic-related pathways including oxidative phosphorylation, the TCA cycle, glycolysis, and gluconeogenesis (Figure 6F; Supplemental File 6). These findings provide a link between mitochondrial structural changes observed with aging and the metabolic basis of age-related diseases. Additional pathway analyses support the notion that Marf KD leads to mitochondrial dysfunction, metabolic defects, and abnormal reactive oxygen species production while also inhibiting cell division processes (Supplemental Figure 7; Supplemental File 7). RNA sequencing also revealed that mitochondrial biogenesis and its upstream transcriptional regulators (e.g., PPARGC1A, Esrra) are generally inhibited by Marf KD and correlate with changes in mitophagy, sphingolipids, mammalian target of rapamycin (mTOR) and DNA synthesis (Figure 6 G, Supplemental Figure 8).

**Figure 6.**
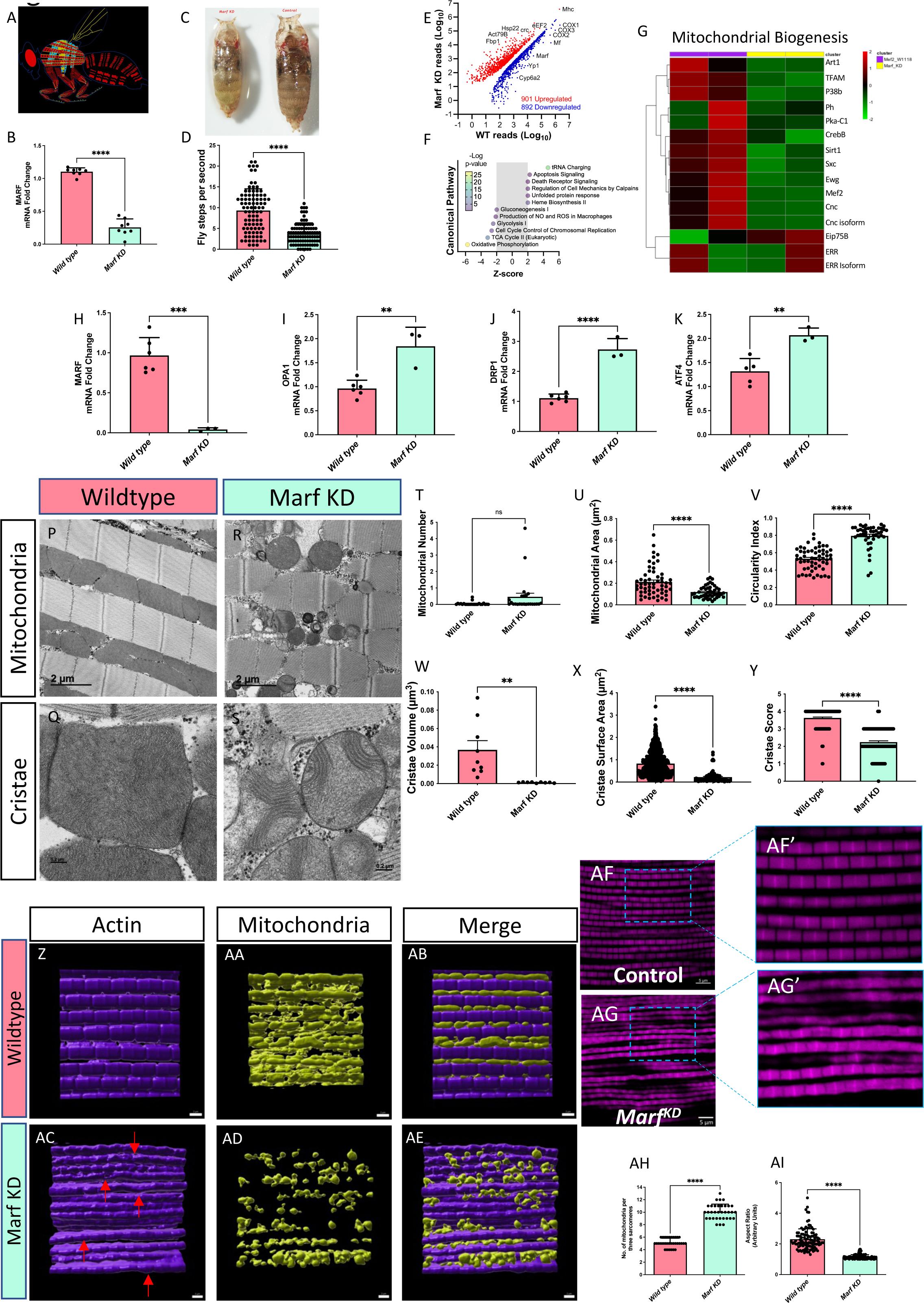
Comparative Analysis of the Impact of Mitochondrial Assembly Regulatory Factor Knockdown (Marf KD) on Mitochondrial Biogenesis and Cellular Features. (A) Schematic representation of the study organism, highlighting specific anatomical regions of flight muscle. (B) Validation of Marf KD through mRNA fold changes, as determined by quantitative PCR (n=8). (C) Visual comparison of wild-type (left) and Marf KD (right) organisms. (D) Fly step quantity changes between wild-type and Marf KD organisms highlight functional differences (n=90). (E) Scatter plot comparing RNA-sequencing reads between wild-type and Marf KD muscles showing differentially expressed genes, with upregulated genes in red and downregulated genes in blue. Select genes are indicated. (F) IPA results for enriched Canonical Pathway terms with an absolute activation Z-score > 2. (G) Heatmap displaying genes related to mitochondrial biogenesis, with gradient colors representing altered expression levels in Marf KD animals compared with controls. The full list of gene names corresponding to FlyBase IDs is available in Supplemental File 3. (H–I) Molecular evaluation of wild-type (n=6) and Marf KD (n=3) organisms according to mRNA fold change, as determined by quantitative PCR of fold changes in (H) Marf, (I) OPA1, (J) DRP1, and (K) ATF4. (P–Q) Transmission electron microscopy images of wild-type flight muscle: (P) longitudinal section and (Q) cross-section. (R–S) Transmission electron microscopy images of Marf KD flight muscle: (R) longitudinal section and (S) cross-section. (T) Quantification of mitochondrial number in the region of interest (n=24), (U) mitochondrial area [n=57 (Wild type) and 45 (Marf KD)], and (V) circularity index in both conditions [n=57 (Wild type) and 45 (Marf KD)]. (W) Quantification of cristae volume (n=9), (X) cristae surface area [n=1089 (Wild type) and 82 (Marf KD)], and (Y) cristae score [n=138 (Wild type) and 120 (Marf KD)], in wild-type and Marf KD mitochondria. (Z-AE) Imaris reconstruction of (Z) actin, (AA) mitochondria, and (AB) merged 3D reconstruction in wildtype and (AC-AE) Marf KD. Red arrows denote bending or curving of actin regions of interest. (AF) Immunofluorescence of actin staining in wildtype and (AG) Marf KD, (AF’-AG’) with specific changes in actin magnified. (AH) Quantitation of the number of mitochondria per sarcomere in *Drosophila* flight muscle (n=∼30) and (AI) aspect ratio (ratio of the major axis to the minor axis; n=∼100). (H–K) Each dot represents an independent experimental run or (T–Y; AH–AI) individual mitochondrion values. Significance was determined with the Mann–Whitney test, with ns, *, **, ***, and **** representing not significant, p ≤ 0.05, p ≤ 0.01, p ≤ 0.001, and p ≤ 0.0001.

Based on these pathway and functional changes, we aimed to understand the changes in mitochondrial dynamics caused by the loss of Marf. We first examined the mRNA levels of several key mitochondrial proteins following Marf silencing. As expected, Marf KD decreased mRNA transcripts of Marf (Figure 6H), with slight variation from validation in Figure 6B. Interestingly, however, OPA1 and DRP1 were both increased (Figures 6I–J), indicating the upregulation of dynamic proteins without a clear preference toward fusion or fission. We also noticed that endoplasmic reticulum (ER) stress is increased with upregulation of ATF4 (Figure 6K) which was confirmed by previous studies showing that MFN2 deficiency causes ER stress (Ngoh et al. 2012). We also look at other ER stress proteins of ATF6 and IRE1 (Supplemental Figures 9A–B), Notably, ER stress can result in MERC formation (Wan et al. 2014). Therefore, we further examined MERC proteins along with GRP75 and VDAC3, which were increased and decreased, respectively, with the loss of Marf (Supplemental Figures 9A–B). Together, these findings suggested multiple changes in dynamics with unclear implications, thus we sought to understand specific structural changes using TEM.

When Marf was knocked out (Figures 6P–S), the mitochondrial number did not change (Figure 6T), which may be because of increases in both DRP1 and OPA1, resulting in increased fusion and fission. Additionally, paralleling the results of DKO condition in myotubes, Marf KD resulted in smaller mitochondria with greater circularity (Figures 6U–V). Significant decreases in cristae volume and surface area were observed, indicating large impairments in mitochondrial oxidative phosphorylation (Figures 6W–X). To further consider cristae structure, we used a metric known as the cristae score, which grades cristae from one to four based on their relative quality and quantity, with four representing “healthy” cristae (Lam et al. 2021; Eisner et al. 2017). The cristae score significantly decreased, consistent with loss of cristae integrity along with mitochondrial structure, which is evolutionarily conserved across both murine-derived myotubes and *Drosophila* (Figures 6T–Y).

Imaris 3D reconstructions of actin (Figure 6Z), mitochondria (Figure 6AA), and merged structures (Figure 6AB) in wildtype and Marf KD *Drosophila* flight muscle show marked differences. Beyond confirming TEM results of smaller and more scattered mitochondria in Marf KD, we show that wildtype actin filaments are straight, whereas Marf KD samples (Figures 6AC-AE) exhibit notable bending or kinking (red arrows). This was further confirmed by immunofluorescence staining (Figures 6AF, AG) and magnified views (Figures 6AF’, AG’) which show that Marf KD results in increased actin disorganization compared to wildtype. This indicates that beyond its role in mitochondrial structure, in *Drosophila*, Marf plays a critical role in maintaining the structural integrity of actin filaments, with implications in cytoskeletal dynamics. Finally, we performed quantification of mitochondrial number per sarcomere (Figure 6AH) and mitochondrial aspect ratio (Figure 6AI) in *Drosophila* flight muscle, which showed both numbers of mitochondria per sarcomere and the aspect ratio in Marf KD was lower, implying that changes in actin structure occur concomitantly with alterations in mitochondrial morphology and distribution.

## Discussion

To our knowledge, studies of mitochondrial changes in human skeletal muscle throughout aging remain limited. Three-dimensional reconstructions of mitochondria showed that structural phenotypes in human skeletal muscle shift to a less complex phenotype with age, concomitantly with the loss of proteins associated with mitochondrial and cristae dynamics. Across disparate cohorts, we observed a decrease in muscle size with age, along with cohorts showing age-related muscle atrophy in human skeletal muscle, which may arise in part due to mitochondrial loss of complexity and concomitant structural remodeling. This finding is further supported by the functional implications of limited exercise endurance among aged humans. Moreover, our data show that mitochondrial structure also rearranges in *Drosophila* with loss of Marf, the ortholog for MFN1 and MFN2. This finding suggests an evolutionarily conserved mechanism both *in vivo* and *in vitro* through which aging results in loss of MFN1 and MFN2, causing a decline in mitochondrial architectural integrity, with exercise serving as a potential therapy to restore mitochondrial structure and associated bioenergetics through increases in mitofusins. While our findings are based on general aging in skeletal muscle, they may carry significant translational implications for the development of therapies for sarcopenia. Beyond this, while exercise is commonly recognized as a treatment for sarcopenia, recent findings also underscore that in mitochondrial-dependent mechanisms regular exercise can largely ameliorate the deleterious effects of aging in skeletal muscle (Grevendonk et al. 2021). Based on these findings, several key promising areas must be further explicated in the future.

Our key finding is that the 3D structures of intermyofibrillar mitochondria show a decrease in complexity and reduced branching patterns in an aged cohort, suggesting structural remodeling caused by the aging process. We also found key genes associated with mitochondria and their contact sites are lost during the aging process. Past studies investigating human skeletal muscle 3D structure have shown that intermyofibrillar mitochondria are distinct from other mitochondrial subpopulations, such as subsarcolemmal mitochondria (Vincent et al. 2019), which are more interconnected than intermyofibrillar mitochondria (Dahl et al. 2015). Thus, investigating whether the structure of subsarcolemmal mitochondria changes during aging may offer insight into how mitochondrial subpopulations differentially respond to sarcopenia. Specifically, another study using FIB-SEM (Marshall, Damo, et al. 2023) showed that the 3D structure of Type I and Type II human skeletal muscle mitochondria also differed, with Type II having lower-volume mitochondria (Dahl et al. 2015). Notably, in sarcopenia, a preferential loss of Type II (fast-twitch) muscle fibers occurs, leading to an increased proportion of Type I (slow-twitch) fibers, for which previous studies have found higher mitochondrial fusion rates (Bellanti et al. 2021). However, our present study did not allow for the differentiation of these fiber types in 3D; therefore, future investigations should consider the differential interplay between mitochondrial structure and exercise across these different fiber types and subpopulations.

The other key finding we noted was that mitochondrial volume increased in aged samples. Since MFN2 is a fusion protein (Chen et al. 2003), this was unexpected, since the import of MFN2 is often discussed in the context of preventing fragmentation, with exercise delaying age-related mitochondrial fragmentation (Campos et al. 2023). However, past studies in skeletal muscle have shown that deletion of *Mfn2* results in impaired electron transport chain complex I activity and mitochondrial swelling, which is caused by osmotic changes (Luo et al. 2021). Within a past study looking at murine skeletal muscle sarcopenia, Leduc-Gaudet and colleagues noticed that subsarcolemmal mitochondria were larger marked by a decreased Mfn2-to-Drp1, as compared to their young counterparts (Leduc-Gaudet et al. 2015). Therefore, when considering the structure-function relationship of mitochondria, the larger mitochondrial volume in aged samples may not confer enhanced bioenergetics. Instead, it could be indicative of impaired function. Swelling, which typically occurs in response to Ca^2+^-overload or oxidative stress, can cause abnormal cristae structure (Shibata et al. 2019), such as that which we observed in MFN2 and Marf KD conditions. Other results in murine cardiac tissue have shown that age-dependent swelling is concomitant with reductions in the ATP production (Rosa et al. 2023). Swelling, accompanied by loss of membrane potential, typically occurs antecedent to the mitochondrial permeability transition pore (mPTP), which can ultimately lead to cell death (Jenkins et al. 2024; Safiulina et al. 2006). Indeed, the uncoupling of mitochondria from Ca2L release units which occurs with age in skeletal muscle (Pietrangelo et al. 2015), may lead to increased vulnerability to mPTP opening, which is a hallmark of murine aging (Cartee et al. 2016). The potential of exercise to mitigate this age-dependent mPTP sensitivity has, in part, led to exercise being proposed as a key mechanism for healthy aging (Cartee et al. 2016; Heo et al. 2018). Particularly, MFN2 has a modulatory effect on calcium owing to its role in mitochondria-ER contacts (Yang et al. 2023), which is suggestive of a role in mitigating mPTP sensitivity. Yet, conflicting findings, mostly in cardiac tissue, show that MFN2 deletion protects against Ca^2+^-overload (Papanicolaou et al. 2012; Chen et al. 2021), while other findings show that MFN2 deletion leads to Ca^2+^-overload in mouse embryo fibroblasts in response to ER stress (Muñoz et al. 2013). It may be that temporary MFN2 activation increases mPTP sensitivity, such as that seen immediately following exercise (Magalhães et al. 2013), while long-term effects of MFN2 of mitochondrial swelling and mPTP sensitivity have longer-term roles in skeletal muscle that remain poorly established.

While we examined vastus lateralis, thigh, and quadricep muscles in this study, an interesting future avenue may be to compare these findings to the 3D structure of biceps and other skeletal muscle regions. We showed that, during aging, the reductions in upper body endurance are not as severe as those observed for lower body endurance. This result may be due to variations in mitochondrial structure remodeling with aging in different regions of skeletal muscle in humans. While skeletal muscle mitochondria evidently impact muscle function, consideration of the wider neuromuscular system (Rygiel et al. 2016) suggests that modulation and changes in neuronal 3D mitochondrial structure may confer increased susceptibility to sarcopenia, which remains poorly elucidated.

One intriguing aspect of our study is the sex-dependent difference in muscle mass loss as individuals age. While male participants showed a significant decrease in thigh CSA, females demonstrated an increase in femur CSA. However, we noted a similar bone-to-thigh ratio for both sexes. Furthermore, there were slightly different changes in endurance with aging. Despite that the literature has referred to sex-specific aging impacts on sarcopenia (Tay et al. 2015), few studies have examined the interplay between these changes and mitochondrial dynamics in muscle tissue. In this study, we combined male and female samples for 3D reconstruction due to the laborious nature of this process, but we generally showed little sex-dependency comparing mitochondria phenotype between male and female samples. Since we did not perform a rigorous analysis to study sex-age interaction effects, future studies may further explore the sex-related changes in the 3D structure of human skeletal muscle mitochondria.

Mitochondrial complexity loss occurs concomitantly with declines in endurance and exercise strength during aging. The restoration of MFN2 levels following exercise suggests mitochondrial structure can be repaired. However, the limitations of *in vivo* studies make it difficult to predict exactly how mitochondrial structure may remodel in response to exercise. Generally, previous *in vivo* studies have suggested that exercise induces the same increase in MFN1 and MFN2 as we observed with our *in vitro* study (Axelrod et al. 2019; Anon n.d.). In our study, we only saw significant increases in MFN2 and OPA1 in old exercising adults, suggesting that exercise may specifically mitigate the age-related loss of MFN2 and OPA1. Future studies must determine how mitochondrial structure remodels immediately following exercise and the long-term benefits of exercise regimens. While this remains difficult *in vivo*, the *in vitro* method for electrical pulse stimulation exercise may facilitate this study and further establish whether exercise alone can prevent mitochondrial remodeling. Previous studies have shown that murine mitochondria subjected to 3-hour acute exercise do not exhibit alterations in size and morphology (Picard et al. 2013), yet few studies have investigated the effects on 3D morphology or differed exercise regimens. Another study applied a chronic 12-week resistance exercise training program, which showed that coupled mitochondrial respiration increased concomitantly with increased muscle strength, but mRNA transcripts of mitochondrial markers of bioenergetics were unchanged (Porter et al. 2015). Other findings have shown that exercise in *C. elegans* with abrogated MFN1 and MFN2 orthologs had impaired physical fitness (Campos et al. 2023). Thus, a promising future avenue is optimizing exercise regimens based on how they change mitochondrial dynamic proteins and associated mitochondrial 3D structure, offering a research-backed avenue to identify the types of exercise that provide the most therapeutic benefits to prevent age-related muscle loss.

Despite MFN2 being a MERC tether protein (Basso et al. 2018; De Brito & Scorrano 2008; Filadi et al. 2015), the loss of Marf induced additional MERC formation. Notably, the role of MFN2 in MERC formation remains controversial; it has been noted as a tether protein in both cultured cells and *in vivo* (Naon et al. 2016; Han et al. 2021; Zaman & Shutt 2022), yet some findings have shown that loss of MFN2 acts to tether MERCs (Cieri et al. 2018; Zaman & Shutt 2022). This finding suggests that MERCs may be formed by the upregulation of other MERC proteins in response to loss of MFN2, including VDAC and PACS2, as a compensatory mechanism during loss of Marf. Similarly, we noticed that while MFN2 was lost in human and murine samples with age, GRP75, IP_3_R3, and VDAC3 were all increased, suggesting greater capacity for MERC formation. However, our qualitative analysis of MERCs in aging was unclear, showing that 3D MERC length may be reduced in aged samples, yet more contacts occur (Figure 2). This suggests an MFN2-mediated loss may still cause MERC widening despite compensatory increases in GRP75, IP_3_R3, and VDAC3 which cause more individual MERCs. Alternatively, we recently showed that loss of OPA1, which occurs in the aging process (Tezze et al. 2017), also causes induction of ER stress and associated MERC tethering (Hinton et al. 2024). Therefore, it remains unclear whether MFN2 loss and OPA1 have pluralistic effects on MERC formation. It is possible that as mitochondria become less complex during aging, MERC tethering changes as a compensatory mechanism because the reduced branching and surface area limit the area for MERCs. Contrary to our findings, previous studies have found that exercise in murine models decreases MERC formation (Merle et al. 2019), potentially through mechanisms involving decreased ER stress (Kim et al. 2017). Thus, it remains unclear whether MERC formation is also involved in alterations in mitochondrial structure in exercise and aging. It also remains unclear whether MFN2 is mediating these pathways in exercising, and future studies must rigorously perform quantitative analysis of MERCs in 2D and 3D EM (Hinton et al. 2023). Therefore, future studies may seek to better elucidate how MERC phenotypes change in aged human skeletal muscle and determine whether these changes are independent of OPA1-mediated alterations.

An important future avenue is better exploring the role of FGF21 in sarcopenia. FGF21, as previously demonstrated, is a universal metabolism regulator that is important for modulating insulin sensitivity (Potthoff 2017). Notably, recent studies have revealed that FGF21 serum levels are increased in individuals with sarcopenia and are directly correlated with loss of muscle strength (Jung et al. 2021; Roh et al. 2021). FGF21’s principal inductor is ER stress, specifically through pathways involving ATF4 (Wan et al. 2014). Notably, ATF4 has also recently arisen as a key modulator of MERCs through OPA1-dependent pathways in skeletal muscle (Hinton et al. 2024). We also showed that ATF4 is increased with Marf KD. This finding suggests that FGF21 may be involved in MERC formation, which may have functional implications in the development of sarcopenia. However, few studies have investigated FGF21-dependent MERC formation. Mitofusins are understood to be required for glucose homeostasis and modulation of insulin sensitivity (Georgiadou et al. 2022; Sebastián et al. 2012). FGF21 may thus provide a mechanistic link through MFN-mediated development of sarcopenia through modulational of MERCs and functional implications of altered glucose metabolism due to mitochondrial structural rearrangement.

Our work also delved into the commonality of exercise and aging as producing similar phenotypes in some cases. Beyond FGF21, we also found that exercise affects IL6 and the immune response, which are observed to be hallmarks of aging (Ershler et al. 1993). While the literature is notably sparse on the molecular pathways that exercise influences to provide its anti-aging benefits, it is possible that immune responses are involved in these changes. This increase in IL-6 that occurs with exercise may occur through the same mechanism in which aging causes IL-6 uptick: ROS generation; however, the exact mechanism remains poorly elucidated (Fischer 2006). Similarly, reduced oxidative stress may concomitantly decrease IL-6 (Lowes et al. 2013). Notably, PGC-1α expression, an important regulator of mitochondrial biogenesis, is linked to both IL-6 and aging, with loss of IL-6 having an inverse effect on PGC-1α levels (Bonda et al. 2017) and causing increased mitochondrial replication (Skuratovskaia et al. 2021). Beyond this, the IL-6 upregulates MFN1 to cause mitochondrial fusion, suggesting a potential mechanism through which exercise stimulates the mitochondrial fusion (Hou et al. 2023). It has been proposed that repeated exercise training can reduce age-related increases in IL-6 (Fischer 2006), suggesting exercise may be useful in modulating the immune response, but how age-dependent changes in IL-6 may contribute to mitochondrial remodeling remains an avenue for greater investigation.

Furthermore, independent from exercise, it is unclear whether MFN2 can be delivered to recapitulate the therapeutic effects of exercise on sarcopenia. Previously, loss of MFN1 and MFN2 was shown to impede exercise performance, suggesting that the age-related loss of endurance we observed is due to MFN2 (Bell et al. 2019). MFN1 and MFN2 also regulate glucose homeostasis through the determination of mtDNA content (Sidarala et al. 2022). Similarly, mtDNA content is decreased following exercise (Puente-Maestu et al. 2011). Given that high circulatory levels of mtDNA are associated with sarcopenia (Fan et al. 2022), this finding offers another potential mechanism through which exercise-mediated restoration of MFN2 protects against sarcopenia, but further investigation into how mtDNA content may alter mitochondrial structure is valuable. Regardless of the specific mechanism, measuring the therapeutic potential of MFN2 is a valuable future avenue. Notably, MFN2 loss was associated with myocardial hypertrophy in cardiac tissue, while gene delivery of an adenoviral vector encoding rat MFN2 proved to successfully protect against myocardial hypertrophy (Yu et al. 2011). However, it is unclear whether these techniques can successfully be applied to humans and if supplementation of MFN2 levels is an effective therapy independent of exercise. Furthermore, past studies have shown that active women do not have elevated MFN2 levels (Drummond et al. 2014), while other studies have shown the opposite, with immediate increases in MFN1 and MFN2 following exercise in athletes (Anon n.d.). These results suggest that further research on the long-term effects on MFN2 protein levels following exercise is necessary.

Taken together, we found evidence that several components related to mitochondrial dynamics, specifically proteins involved in mitochondrial fusion and fission, as well as MERCs, were altered during the aging process. This underscores our previous findings (Vue, Garza-Lopez, et al. 2023; Vue, Neikirk, et al. 2023), which show that, beyond only fusion and fission dynamics, MERCs and mitochondrial 3D structure must all be considered in the aging process. Our study remains limited by having cohorts defined as “old” cut-off of 50 years old, with “young” samples potentially displaying heterogeneity characteristic of middle-aged participants. Additionally, we are limited by the necessity of several cohorts that correlatively but not causatively show changes in MRI parameters, mitochondria structure, exercise ability, and plasma immune factors. While we were unable to prove that these changes concomitantly occur in old patients with sarcopenia, this study suggests the simultaneous study of these factors in a human model remains important in future studies. Our cross-species analysis using *Drosophila* models provided compelling evidence that the mechanisms we observed are evolutionarily conserved. While we are not the first to show that exercise increases MFN2 protein levels opposite to losses caused by aging (Koltai et al. 2012; Cartoni et al. 2005), the structural impacts of these changes have remained ambiguous. While our findings further our understanding of age-dependent mitochondrial structure, the physiological implications of reduced mitochondrial complexity, the specific molecular mechanisms through which exercise confers its benefits, and the evolutionary conservation of these mechanisms remain new avenues for therapeutic interventions to counteract the deleterious effects of aging.

## Data Sharing and Open Access

All data are available upon request to the corresponding author.

## Author Contributions

E.S., Z.V., P.K., A.M., L.V., E.G.L., K.N., D.S. drafted the manuscript and performed experiments. D.M. D.D.H., R.R., J.S., M.M., S.T.A., I.H., S.M., C.W., A.W., C.W., S.M.D., J.A.G., A.K., B.G., E.H.M.D., A.K., F.S., M.B. contributed to the study design and data analysis. M.R.M., M.A.P., A.C., S.A.M., V.E., B.C.M., A.H. provided data curation and project administration. All authors contributed to the design of the study, data interpretation, and manuscript revision.

## Videos

Videos 1-9: Representative 3D reconstruction of mitochondria from young cases (Case #1: Video 1; Case #2: Video 2; Case #3: Video 3; Case #4: Video 4; Case #5: Video 5) and old cases (Case #2: Video 7; Case #3: Video 8; Case #4: Video 9) of human skeletal muscle.

Video 10-11: Examples of exercises performed by Cohort #3: chair stand test as a proxy for lower body endurance (Video 10) and arm curl test as a proxy for upper body endurance (Video 11).

Videos 12–15: Behavior and motor function of control *Drosophila* (Videos 12–13) and Marf knockdown organisms (Videos 14–15)

## Supplemental Figures

**Supplemental Figure 1: Schema of 5 cohorts used to analyze age-related changes in skeletal muscle.**

**Supplemental Figure 2: Sex-dependent Differences in Magnetic Resonance Imaging Measurements.** (A) Grouped measurements of thigh, femur, tibia, and total calf muscle cross-sectional area (CSA) in males and females. (B) Scatter box plot detailing thigh CSA, (C) femur CSA, (D) tibia CSA, (E) and total calf muscle CSA across young males, young females, older males, and older females. Intra- and inter-sex-dependent differences during aging are compared. (A) Multiple Mann–Whitney tests with the two-stage step-up method of Benjamini, Krieger, and Yekutieli were used to correct for the false discovery rate. (B–E) Intergroup comparisons were performed using one-way ANOVA with Tukey’s multiple comparisons test *post hoc*. Statistical significance is denoted as ns (not significant), *p < 0.05, **p < 0.01, ***p < 0.001, ****p < 0.0001.

**Supplemental Figure 3: Heterogeneity in Mitochondrial Quantification Across Patients.** (A) Representative images of 5 young cases (mitochondrial number varies; Case #1: n = 253; Case #2: n = 250; Case #3: n = 250; Case #4: n = 252; Case #5: n = 253; total mitochondria surveyed across young cohort: n = 1258) and 4 old cases (mitochondrial number varies; Case #1: n = 254; Case #2: n = 250; Case #3: n = 250; Case #4: n = 250; total mitochondria surveyed across old cohort: n = 1004). Distribution of mitochondria for patient heterogeneity in (B) mitochondrial volume, (C) surface area, (D) perimeter, (E) sphericity, and (F) complexity index in young and old human skeletal muscle.

**Supplemental Figure 4: Protocols for performing exercises.** (A) Walking where participants were tasked with walking the maximum distance in a course designed with 45.72 meters for a 6-minute timer. (B) Grip strength was measured in each arm through participants’ maximum grip with their forearms at a 90° angle. (C) Localized muscle endurance (LME) of the lower body was measured with participants seated on a chair with their back against a wall for greater stability, and they performed the maximum number of complete raises for a 30-second time period. (D) LME of the upper body was assessed through an adapted method in which participants were seated on a chair while performing the maximum number of unilateral elbow flexions for a 30-second time period with a 4kg (men) or 2kg (women) weight.

**Supplemental Figure 5: Differences in Exercise Parameters Between Young and Old Humans Across Both Sexes.** (A) Chart representing various parameters (weight, height, body mass index (BMI), walking distance, VO_2_max, right and left grip strength, and muscle endurance of the upper and lower body) comparing young and old individuals. Blue bars represent young individuals, and purple bars represent older individuals. (B) Scatter box plot detailing weight distribution across young males, young females, older males, and older females for comparison of intra- and inter-sex-dependent differences in aging. (C) Scatter box plot illustrating the distribution of BMI values, (D) walking distances (in meters), and (E) VO_2_max values during a walking test among the same groups. (F–G) Scatter box plots for grip strength in kg: (F) left grip strength and (G) right grip strength across the four demographic groups. (H–I) Scatter box plots representing localized muscle endurance (H) of the lower body and (I) the upper body across young males, young females, older males, and older females. (A) Multiple Mann–Whitney tests with the two-stage step-up method of Benjamini, Krieger, and Yekutieli were used to correct for the false discovery rate. (B–I) Intergroup comparisons were performed using one-way ANOVA with Tukey’s multiple comparisons test *post hoc*. Statistical significance is denoted as ns (not significant), *p < 0.05, **p < 0.01, ***p < 0.001, ****p < 0.0001.

**Supplemental Figure 6: Full Screening of Various Parameters or Biomolecules in the Blood of Older and Young Individuals.** The chart displays the full extent of parameters considered when comparing young and older individuals, with a mixture of males and females. Multiple Mann–Whitney tests with the two-stage step-up method of Benjamini, Krieger, and Yekutieli were used to correct for the false discovery rate. Statistical significance is denoted as ns (not significant) or * (significant).

**Supplementary Figure 7: RNA-sequencing Pathway Analyses Following Marf Knockdown (Marf KD)** (A and B) Bubble plots of Gene Set Enrichment Analysis (GSEA) showing enriched cellular components (A) and biological processes (B) impacted by Marf KD compared with the wild-type condition. (C-E) Bubble plots of Ingenuity Pathway Analysis (IPA) results for terms annotated for Diseases or Functions (C), Upstream Regulators (D), and Canonical Pathways (E). Data for (E) is the same as in main Figure 6E but is expanded here to include additional terms having an absolute activation Z-score > 1.5. For all panels, enrichment and activation Z-scores between −2 and +2 are indicated by a gray box and are considered insignificant. The color of each term symbol reflects the -Log p-value or false discovery rate (FDR) as indicated by the color scale.

**Supplementary Figure 8: Heatmap Analysis of Pathways Altered Following Marf Knockdown (Marf KD).** Heatmaps of altered expression of proteins associated with (A) mitophagy, (B) mammalian target of rapamycin (mTOR), (C) sphingolipid signaling, and (D) DNA synthesis. The full list of gene names corresponding to FlyBase IDs is available in Supplemental File 6. The color scale on the right side represents expression values, with red indicating upregulation and green indicating downregulation.

**Supplementary Figure 9: Molecular evaluation of wild-type and Marf KD organisms according to mRNA fold change, as determined by quantitative PCR (qPCR)**. qPCR of endoplasmic reticulum stress proteins: (A) ATF6 and (B) IREI1. qPCR of mitochondria– endoplasmic reticulum contact site proteins: (C) GRP75 and (D) VDAC.

## Supplemental Files

Supplemental File 1. Full parameters of Cohort #1, used for thigh magnetic resonance imaging, including patient age.

Supplemental File 2. Full parameters of Cohort #2, used for calf magnetic resonance imaging, including patient age.

Supplemental File 3. Full parameters of Cohort #3, used for imaging of 3D reconstruction, including patient age.

Supplemental File 4. Full parameters of Cohort #4, which was subjected to exercise, including patient age and weight.

Supplemental File 5. Full parameters of Cohort #5, which had blood plasma measured, including patient age and weight.

Supplemental File 6. Orthologs of FlyBase ID names and fly gene names.

Supplemental File 7. Significant differentially expressed genes with GSEA and IPA analysis results.

