## Supplementary material for "3D Mitochondrial Structure in Aging Human Skeletal Muscle: Insights into MFN-2 Mediated Changes": Figures

**Authors and Affiliations:**

Estevão Scudese^1,2,3^*, Zer Vue^1^*, Prassana Katti^4,5*^, Andrea G. Marshall^1^, Mert Demirci^6^, Larry Vang^1^, Edgar Garza López^7^, Kit Neikirk^1^, Bryanna Shao^1^, Han Le^1^, Dominique Stephens^1^, Duane D. Hall^7^, Rahmati Rostami^8^, Taylor Rodman^1^, Kinuthia Kabugi^1^, Chanel Harris^1^, Jian-qiang Shao^9^, Margaret Mungai^1^, Salma T. AshShareef^7^, Innes Hicsasmaz^7^, Sasha Manus^1^, Celestine Wanjalla^10^, Aaron Whiteside^1,11^, Revathi Dasari^4^, Clintoria Williams^11^, Steven M. Damo^12^, Jennifer A. Gaddy^6,13^, Brian Glancy^5,14^, Estélio Henrique Martin Dantas^2,15,16,17,18^, André Kinder^19^, Ashlesha Kadam^20^, Dhanendra Tomar^20^, Fabiana Scartoni^2^, Matheus Baffi^3^, Melanie R. McReynolds^21^, Mark A. Phillips^22^, Anthonya Cooper^23^, Sandra A. Murray^23^, Anita M. Quintana^24^, Vernat Exil^25^, Annet Kirabo^6^, Bret C. Mobley^26^#, Antentor Hinton^1^#

^1^ Department of Molecular Physiology and Biophysics, Vanderbilt University, Nashville, TN, 37232, USA

^2^ Laboratory of Biosciences of Human Motricity (LABIMH) of the Federal University of State of Rio de Janeiro (UNIRIO), Rio de Janeiro, Brazil

^3^ Sport Sciences and Exercise Laboratory (LaCEE), Catholic University of Petrópolis (UCP), Brazil

^4^ Department of Biology, Indian Institute of Science Education and Research (IISER) Tirupati, AP, 517619, India

^5^ National Heart, Lung, and Blood Institute, National Institutes of Health, Bethesda, MD, 20892, USA

^6^ Department of Medicine, Division of Nephrology and Hypertension, Vanderbilt University Medical Center, Nashville, Tennessee, USA

^7^ Department of Internal Medicine, University of Iowa, Iowa City, IA, 52242, USA

^8^ Department of Genetic Medicine, Joan & Sanford I. Weill Medical College of Cornell University, New York, NY, 10065, USA

^9^ Central Microscopy Research Facility, Iowa City, IA 52242, USA

^10^Division of Infection Diseases, Department of Medicine, Vanderbilt University Medical Center, Nashville, TN, 37232, USA

^11^ Department of Neuroscience, Cell Biology and Physiology, Wright State University, Dayton, OH, 45435, USA

^12^ Department of Life and Physical Sciences, Fisk University, Nashville, TN, 37208, USA

^13^Tennessee Valley Healthcare Systems, U.S. Department of Veterans Affairs, Nashville, TN, 37212, USA

^14^ NIAMS, NIH, Bethesda, MD, 20892, USA

^15^ Doctor’s Degree Program in Nursing and Biosciences - PpgEnfBio, Federal University of the State of Rio de Janeiro - UNIRIO, Rio de Janeiro, RJ, Brazil

^16^ Laboratory of Human Motricity Biosciences - LABIMH, Federal University of the State of Rio de Janeiro - UNIRIO, RJ, Brazil

^17^ Brazilian Paralympic Academy – APB

^18^ Doctor’s Degree Program in Health and Environment - PSA, Tiradentes University - UNIT, Aracaju, SE, Brazil

^19^ Artur Sá Earp Neto University Center - UNIFASE-FMP, Petrópolis Medical School, Brazil

^20^ Department of Internal Medicine, Section of Cardiovascular Medicine, Wake Forest University School of Medicine, Winston-Salem, NC 27157 USA

^21^ Department of Biochemistry and Molecular Biology, The Huck Institute of the Life Sciences, Pennsylvania State University, State College, PA, 16801, USA

^22^ Department of Integrative Biology, Oregon State University, Corvallis, OR, 97331, USA

^23^ Department of Cell Biology, School of Medicine, University of Pittsburgh, Pittsburgh, PA, 15260, USA^24^

^24^ Department of Biological Sciences, Border Biomedical Research Center, The University of Texas at El Paso, El Paso, Texas, USA

^25^ Department of Pediatrics, Div. of Cardiology, St. Louis University School of Medicine, St. Louis, MO, 63104, USA

^26^ Department of Pathology, Vanderbilt University Medical Center, Nashville, TN, 37232, USA

*These authors share co-first authorship.

#These authors share senior authorship.

Corresponding Author:

Antentor Hinton

Department of Molecular Physiology and Biophysics Vanderbilt University 319-383-3095

Keywords: Mitochondria, Aging, Exercise, Human Skeletal Muscle, 3D Reconstruction, MFN2

Supplemental Figures:

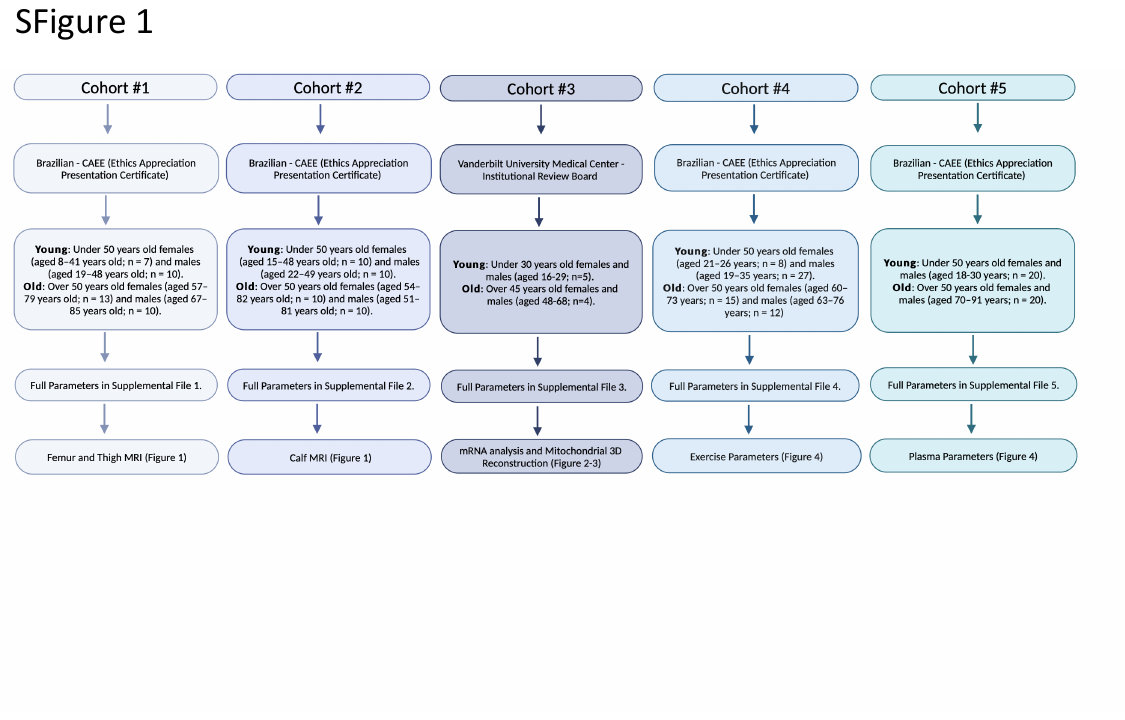

Supplemental Figure 1: Schema of 5 cohorts used to analyze age-related changes in skeletal muscle.

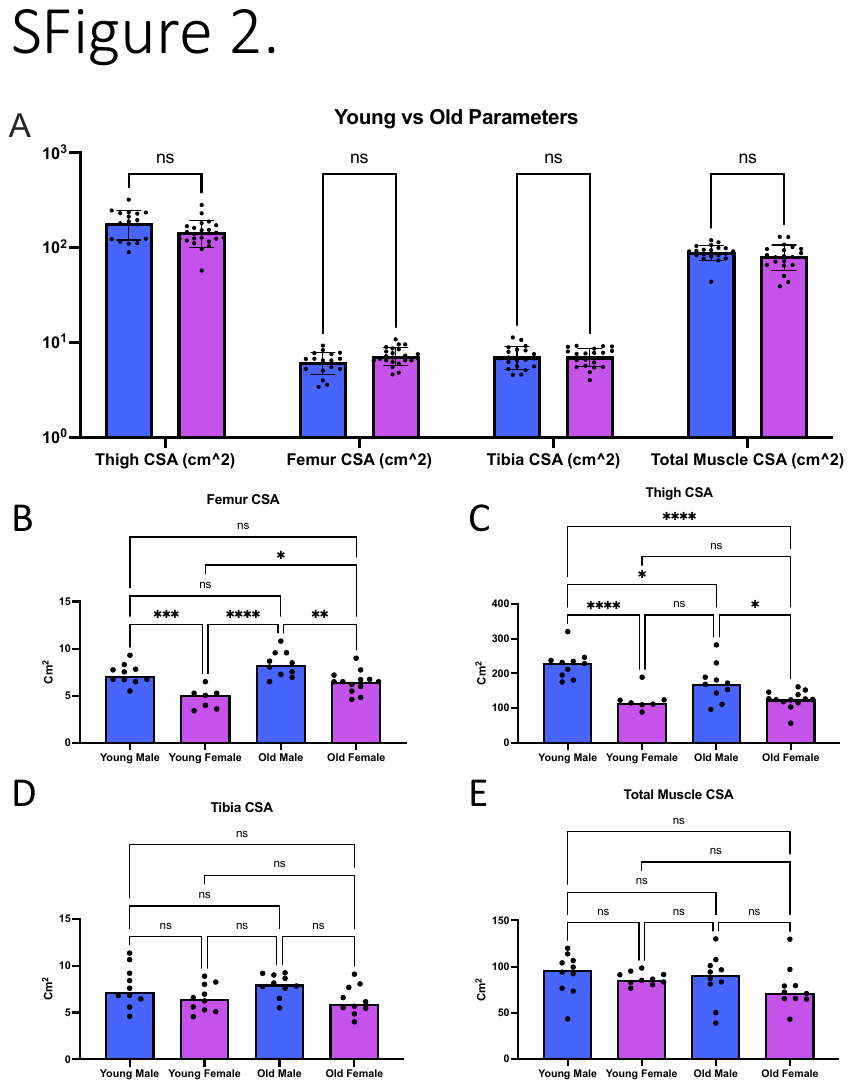

Supplemental Figure 2: Sex-dependent Differences in Magnetic Resonance Imaging Measurements. (A) Grouped measurements of thigh, femur, tibia, and total calf muscle cross-sectional area (CSA) in males and females. (B) Scatter box plot detailing thigh CSA, (C) femur CSA, (D) tibia CSA, (E) and total calf muscle CSA across young males, young females, older males, and older females. Intra- and inter-sex-dependent differences during aging are compared. (A) Multiple Mann–Whitney tests with the two-stage step-up method of Benjamini, Krieger, and Yekutieli were used to correct for the false discovery rate. (B–E) Intergroup comparisons were performed using one-way ANOVA with Tukey's multiple comparisons test *post hoc*. Statistical significance is denoted as ns (not significant), *p < 0.05, **p < 0.01, ***p < 0.001, ****p < 0.0001.

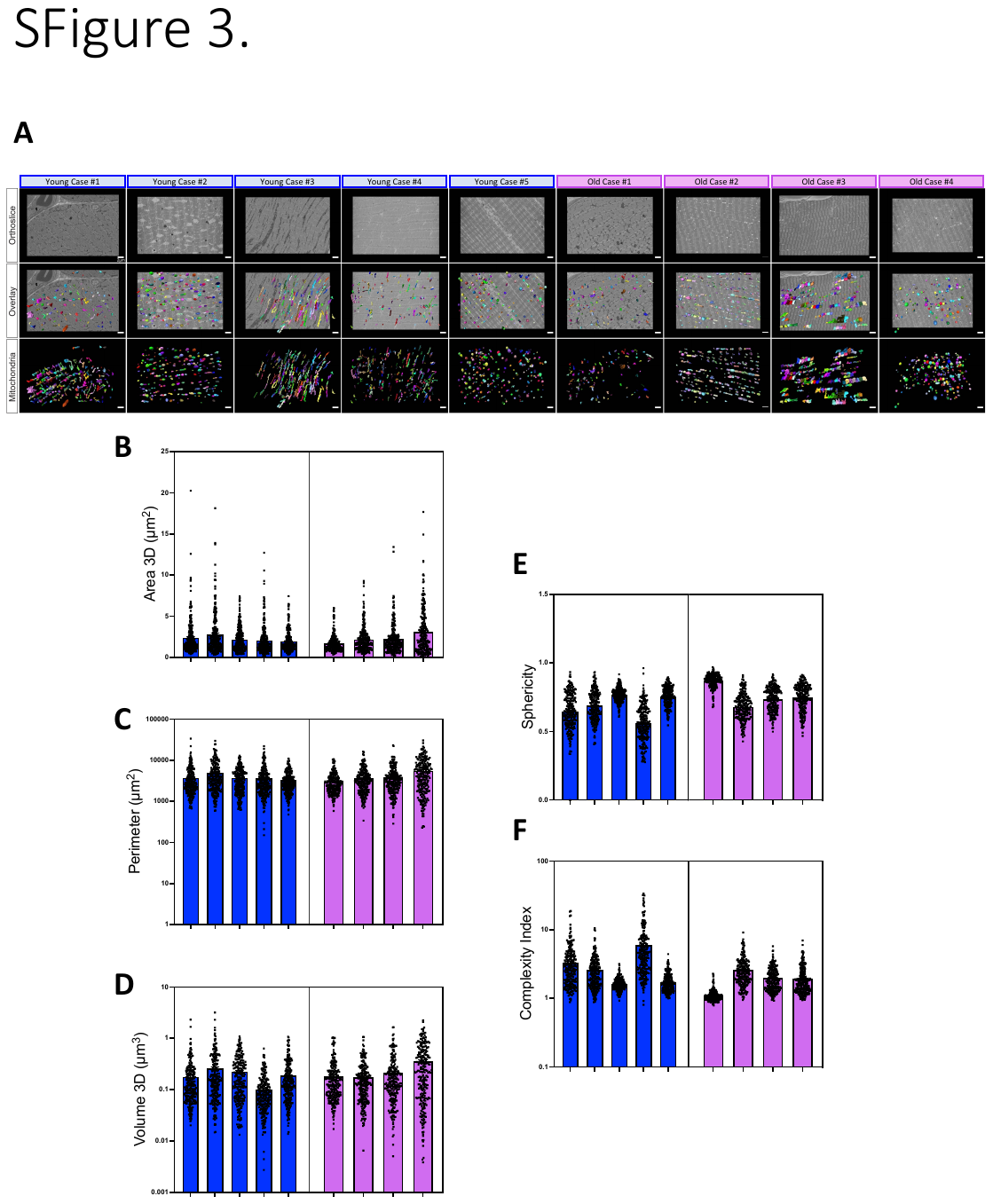

Supplemental Figure 3: Heterogeneity in Mitochondrial Quantification Across Patients. (A) Representative images of 5 young cases (mitochondrial number varies; Case #1: n = 253; Case #2: n = 250; Case #3: n = 250; Case #4: n = 252; Case #5: n = 253; total mitochondria surveyed across young cohort: n = 1258) and 4 old cases (mitochondrial number varies; Case #1: n = 254; Case #2: n = 250; Case #3: n = 250; Case #4: n = 250; total mitochondria surveyed across old cohort: n = 1004). Distribution of mitochondria for patient heterogeneity in (B) mitochondrial volume, (C) surface area, (D) perimeter, (E) sphericity, and (F) complexity index in young and old human skeletal muscle.

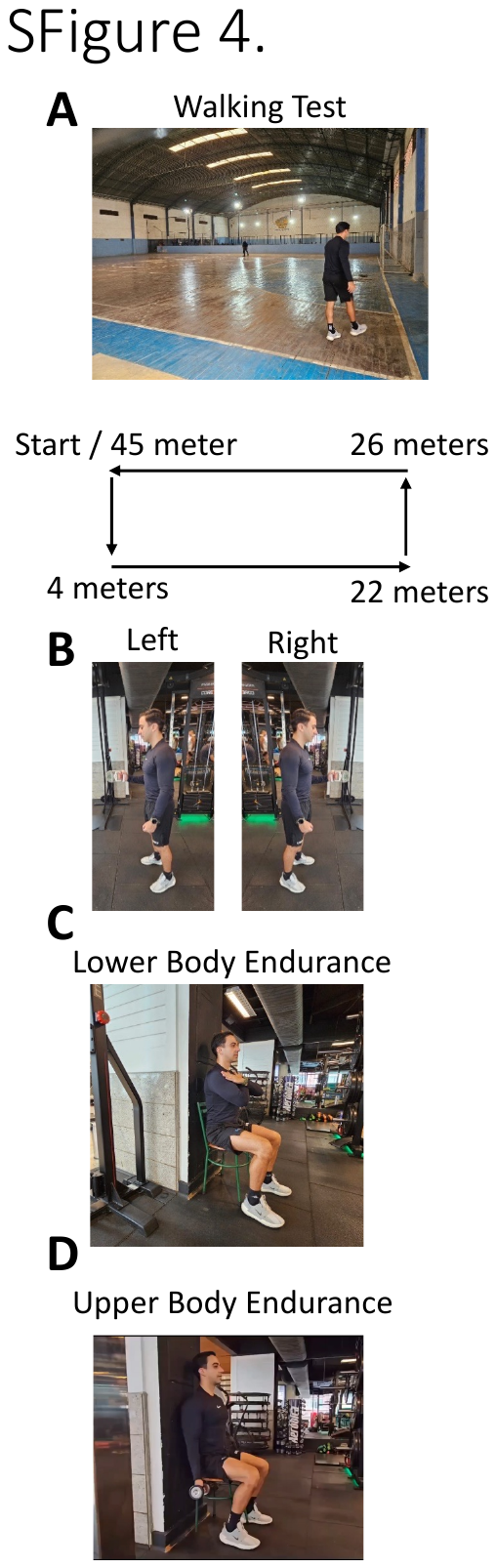

Supplemental Figure 4: Protocols for performing exercises. (A) Walking where participants were tasked with walking the maximum distance in a course designed with 45.72 meters for a 6-minute timer. (B) Grip strength was measured in each arm through participants' maximum grip with their forearms at a 90º angle. (C) Localized muscle endurance (LME) of the lower body was measured with participants seated on a chair with their back against a wall for greater stability, and they performed the maximum number of complete raises for a 30-second time period. (D) LME of the upper body was assessed through an adapted method in which participants were seated on a chair while performing the maximum number of unilateral elbow flexions for a 30-second time period with a 4kg (men) or 2kg (women) weight.

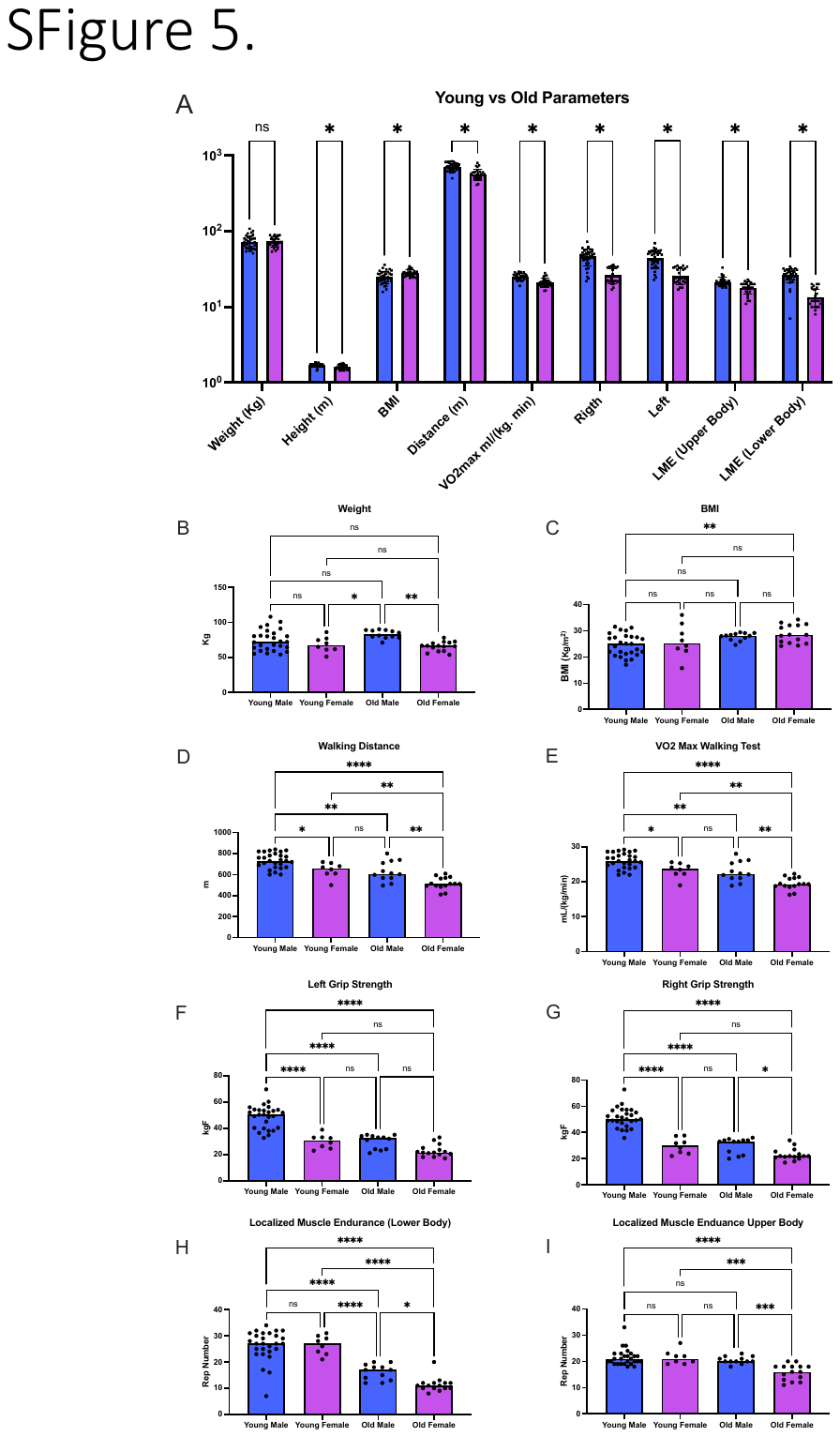

Supplemental Figure 5: Differences in Exercise Parameters Between Young and Old Humans Across Both Sexes. (A) Chart representing various parameters (weight, height, body mass index (BMI), walking distance, VO_2_max, right and left grip strength, and muscle endurance of the upper and lower body) comparing young and old individuals. Blue bars represent young individuals, and purple bars represent older individuals. (B) Scatter box plot detailing weight distribution across young males, young females, older males, and older females for comparison of intra- and inter-sex-dependent differences in aging. (C) Scatter box plot illustrating the distribution of BMI values, (D) walking distances (in meters), and (E) VO_2_max values during a walking test among the same groups. (F–G) Scatter box plots for grip strength in kg: (F) left grip strength and (G) right grip strength across the four demographic groups. (H–I) Scatter box plots representing localized muscle endurance (H) of the lower body and (I) the upper body across young males, young females, older males, and older females. (A) Multiple Mann–Whitney tests with the two-stage step-up method of Benjamini, Krieger, and Yekutieli were used to correct for the false discovery rate. (B–I) Intergroup comparisons were performed using one-way ANOVA with Tukey's multiple comparisons test *post hoc*. Statistical significance is denoted as ns (not significant), *p < 0.05, **p < 0.01, ***p < 0.001, ****p < 0.0001.

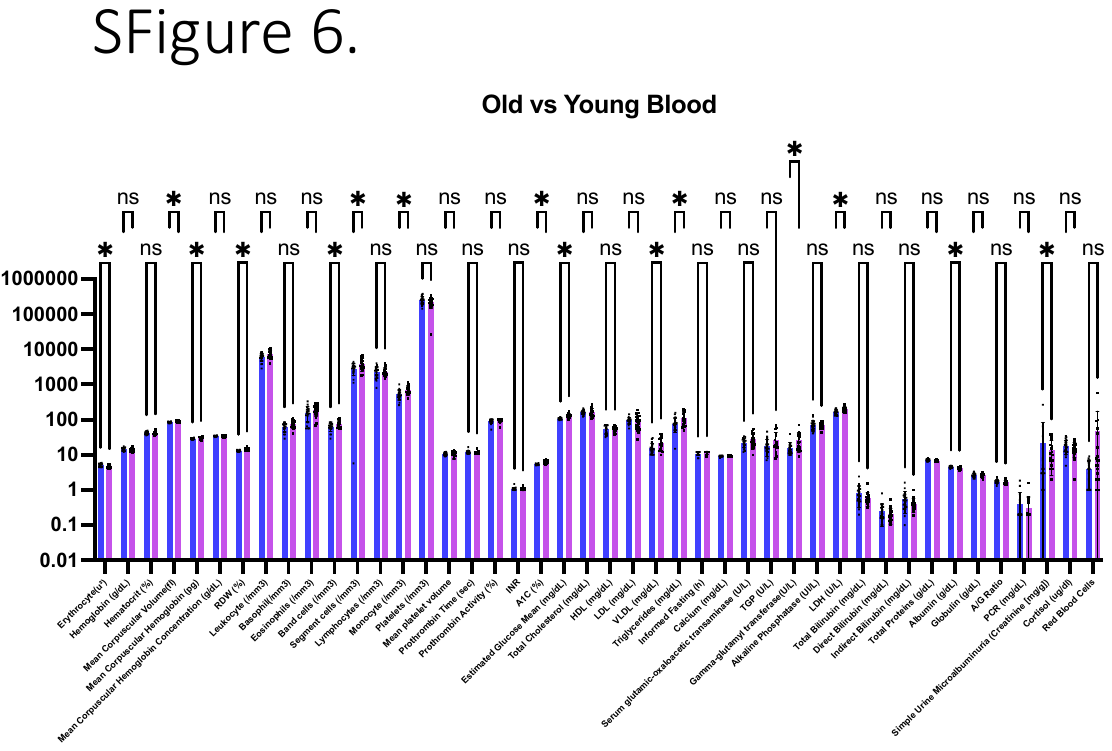

Supplemental Figure 6: Full Screening of Various Parameters or Biomolecules in the Blood of Older and Young Individuals. The chart displays the full extent of parameters considered when comparing young and older individuals, with a mixture of males and females. Multiple Mann–Whitney tests with the two-stage step-up method of Benjamini, Krieger, and Yekutieli were used to correct for the false discovery rate. Statistical significance is denoted as ns (not significant) or * (significant).

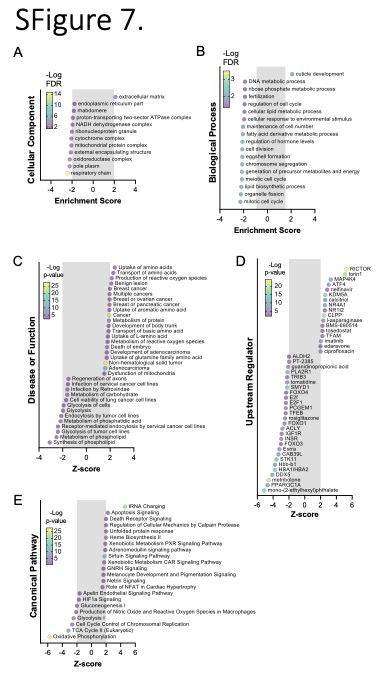

Supplementary Figure 7: RNA-sequencing Pathway Analyses Following Marf Knockdown (Marf KD) (A and B) Bubble plots of Gene Set Enrichment Analysis (GSEA) showing enriched cellular components (A) and biological processes (B) impacted by Marf KD compared with the wild-type condition. (C-E) Bubble plots of Ingenuity Pathway Analysis (IPA) results for terms annotated for Diseases or Functions (C), Upstream Regulators (D), and Canonical Pathways (E). Data for (E) is the same as in main Figure 6E but is expanded here to include additional terms having an absolute activation Z-score > 1.5. For all panels, enrichment and activation Z-scores between -2 and +2 are indicated by a gray box and are considered insignificant. The color of each term symbol reflects the -Log p-value or false discovery rate (FDR) as indicated by the color scale.

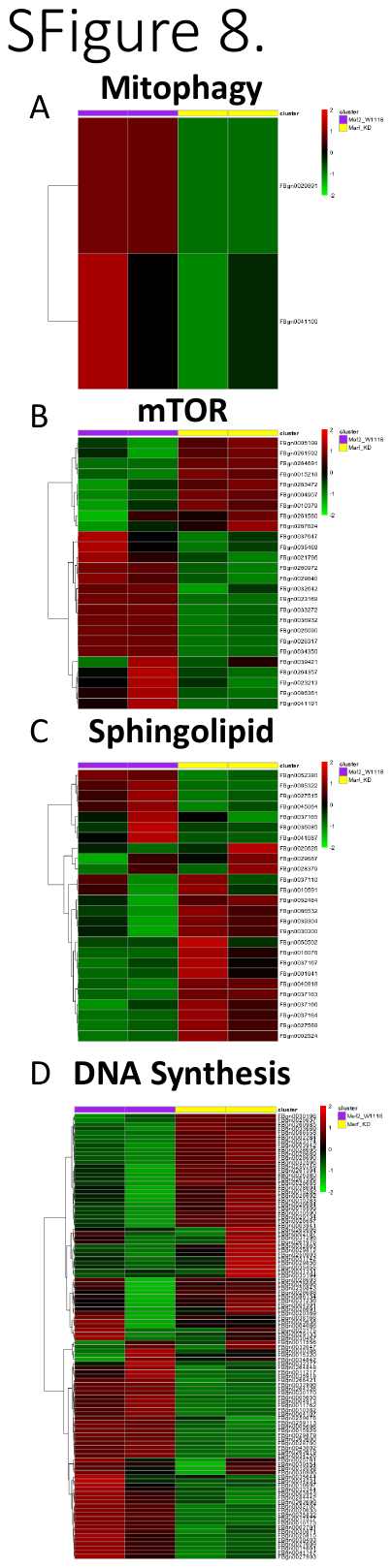

Supplementary Figure 8: Heatmap Analysis of Pathways Altered Following Marf Knockdown (Marf KD). Heatmaps of altered expression of proteins associated with (A) mitophagy, (B) mammalian target of rapamycin (mTOR), (C) sphingolipid signaling, and (D) DNA synthesis. The full list of gene names corresponding to FlyBase IDs is available in Supplemental File 6. The color scale on the right side represents expression values, with red indicating upregulation and green indicating downregulation.

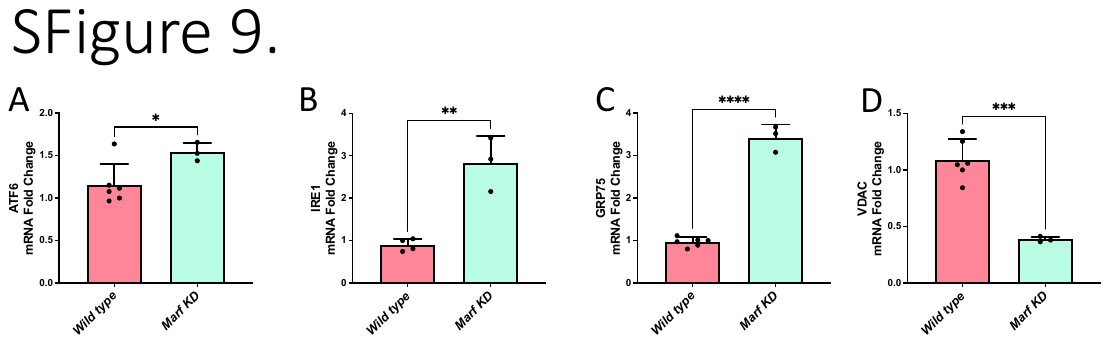

Supplementary Figure 9: Molecular evaluation of wild-type and MARF KD organisms according to mRNA fold change, as determined by quantitative PCR (qPCR). qPCR of endoplasmic reticulum stress proteins: (A) ATF6 and (B) IREI1. qPCR of mitochondria–endoplasmic reticulum contact site proteins: (C) GRP75 and (D) VDAC.

Supplemental Files:

Supplemental File 1. Full parameters of Cohort #1, used for thigh magnetic resonance imaging, including patient age.

Supplemental File 2. Full parameters of Cohort #2, used for calf magnetic resonance imaging, including patient age.

Supplemental File 3. Full parameters of Cohort #3, used for imaging of 3D reconstruction, including patient age.

Supplemental File 4. Full parameters of Cohort #4, which was subjected to exercise, including patient age and weight.

Supplemental File 5. Full parameters of Cohort #5, which had blood plasma measured, including patient age and weight.

Supplemental File 6. Orthologs of FlyBase ID names and fly gene names.

Supplemental File 7. Significant differentially expressed genes with GSEA and IPA analysis results.

Videos:

Videos 1-9: Representative 3D reconstruction of mitochondria from young cases (Case #1: Video 1; Case #2: Video 2; Case #3: Video 3; Case #4: Video 4; Case #5: Video 5) and old cases (Case #2: Video 7; Case #3: Video 8; Case #4: Video 9) of human skeletal muscle.

Video 10-11: Examples of exercises performed by Cohort #3: chair stand test as a proxy for lower body endurance (Video 10) and arm curl test as a proxy for upper body endurance (Video 11).

Videos 12–15: Behavior and motor function of control *Drosophila* (Videos 12–13) and Marf knockdown organisms (Videos 14–15)
